# Modeling Allosteric Mechanisms of Eukaryotic Type II Topoisomerases

**DOI:** 10.1101/2023.08.02.551689

**Authors:** Stefania Evoli, Nilusha L. Kariyawasam, Karin C. Nitiss, John L. Nitiss, Jeff Wereszczynski

**Affiliations:** Department of Physics, Illinois Institute of Technology, Chicago, USA; Center for Molecular Study of Condensed Soft Matter, Illinois Institute of Technology, Chicago, USA; Pharmaceutical Sciences Department, University of Illinois at Chicago, Rockford, IL; Department of Biology, Illinois Institute of Technology, Chicago, USA

## Abstract

Type II topoisomerases (TopoIIs) are ubiquitous enzymes that are involved in crucial nuclear processes such as genome organization, chromosome segregation, and other DNA metabolic processes. These enzymes function as large, homodimeric complexes that undergo a complex cycle of binding and hydrolysis of two ATP molecules in their ATPase domains, which regulates the capture and passage of one DNA double-helix through a second, cleaved DNA molecule. This process requires the transmission of information about the state of the bound nucleotide over vast ranges in the TopoII complex. How this information is transmitted at the molecular level to regulate TopoII functions and how protein substitutions disrupt these mechanisms remains largely unknown. Here, we employed extensive microsecond scale molecular dynamics simulations of the yeast TopoII enzyme in multiple nucleotide-bound states and with amino acid substitutions near both the N- and C-terminals of the complex. Simulation results indicate that the ATPase domains are remarkably flexible on the sub-microsecond timescale and that these dynamics are modulated by the identity of the bound nucleotides and both local and distant amino acid substitutions. Network analyses point towards specific allosteric networks that transmit information about the hydrolysis cycle throughout the complex, which include residues in both the protein and the bound DNA molecule. Amino acid substitutions weaken many of these pathways. Together, our results provide molecular-level details on how the TopoII catalytic cycle is controlled through nucleotide binding and hydrolysis and how mutations may disrupt this process.

**SIGNIFICANCE:** TypeII Topoisomerases (TopoIIs) are essential and ubiquitous for maintaining DNA topology in the nucleus. The mechanisms by which information about the nucleotide-binding state is transmitted from the ATPase domains throughout the TopoII complex remain poorly understood. We used microsecond timescale molecular dynamics simulations to probe the allosteric mechanisms underlying TopoII function. Results indicate remarkable flexibility of the ATPase domains on this timescale which is modulated by nucleotide binding and local and distant amino acid substitutions. Furthermore, we mapped the allosteric networks linking the ATPase and DNA cleavage domains, and connected them to the ATPase domain dynamics. Our findings provide molecular-level insights into how nucleotide binding and hydrolysis regulate the TopoII catalytic cycle and how mutations can disrupt these processes.

## INTRODUCTION

Type II topoisomerases (TopoIIs) catalyze changes in DNA topology and are critical for all organisms with double stranded DNA (1). These ubiquitous enzymes play an essential role in resolving topological problems created during DNA transcription and replication, making them critical in genome organization, chromosome segregation, and other DNA metabolic processes. (2–4). The TopoIIα isoform is essential in cell division and plays a key role in maintaining the structure of mitotic chromatin (5, 6). The TopoIIβ isoform plays key roles in transcription in a variety of tissues and is particularly important in neuronal cells (2, 7). The importance of topoisomerase function in replicating cells has made these enzymes prime targets in the development of antibacterial and chemotherapuetic compounds. (8, 9).

Eukaryotic TopoII enzymes function as large, homodimeric complexes.(1) They remodel DNA through a multistep process that results in the passage of a complete double stranded DNA molecule, the T-segment DNA (tDNA), through a transient double-stranded break in a second DNA molecule, the G-segment DNA (gDNA).(10) The structure of the *Saccharomyces cerevisiae* TopoII complex is a multidomain architecture that can be divided into three major regions (Figure 1)(11). The ATPase domains are built from the GHKL (named for gyrase, HSP90, histidine kinase, and MutL) and transducer domains and contain an ATP binding site that is the location of nucleotide binding and hydrolysis. The DNA-binding/cleavage domain contains the TOPRIM (topoisomerase-primase), winged helix, tower, and coiled-coil domains, and is the location of gDNA binding and cleavage. The C-terminal domain is an unfolded region that has yet to be resolved.

**Figure 1.**
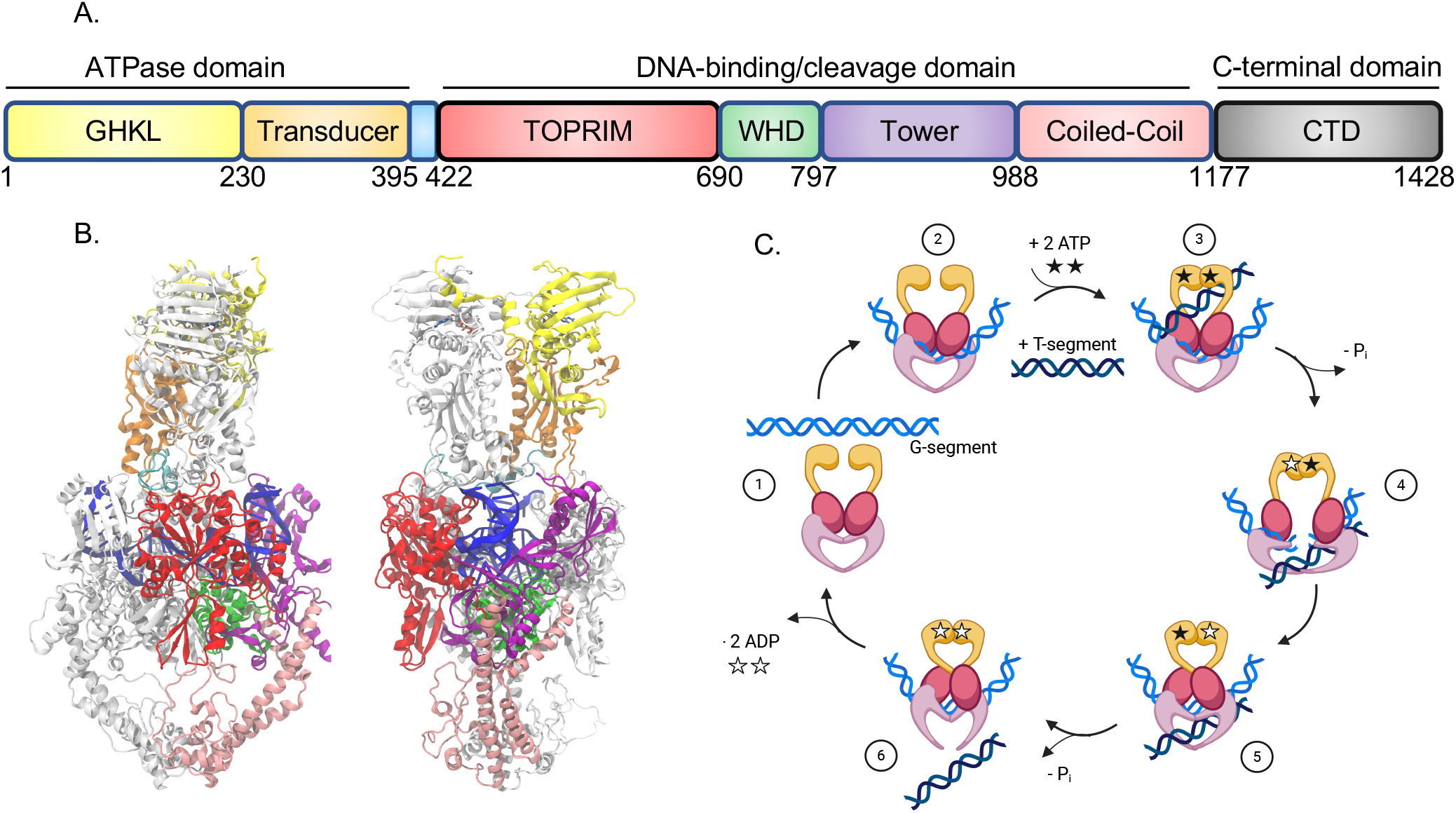
Structure of yeast TopoII when bound to ATP and gDNA with closed N-gate, DNA-gate, and C-gate. (A) Domain arrangement of the ATPase (GHKL and Transducer), DNA-binding/cleavage (TOPRIM, winged helix domain (WHD), tower, and coiled-coil), and C-terminal domains. (B). Crystal structure of yeast TopoII homodimer with one monomer colored according to the domain architecture in (A) and the other colored grey. The C-terminal domain is not included as it is disordered and not resolved in the crystal structure, which was reported by Schmidt *et al*.(11). (C). The topoII catalytic cycle consists of a series of steps including capture of the G-segment DNA, closing of the N-gate upon ATP and T-segment DNA binding, opening of the DNA-gate and passage of the T-segment through the G-segment, and then opening of the C-gate and release of the T-segment.

The working model of the TopoII catalytic cycle requires this complex to function as if it has three “gates” (Figure 1C).(11, 12) Initially, no nucleotides are bound to the ATPase domains, and these domains are separated from one another to form an open “N-gate” structure. Before the N-gate can close, ATP binds to each of the two GHKL domains, and a tDNA segment is captured by the ATPase domains. The disposition of the tDNA segment after capture but prior to DNA cleavage remains poorly understood, since the crystal structure of yeast TopoII cannot obviously accommodate a double stranded DNA unless DNA cleavage has occurred and the DNA-gate has opened. DNA cleavage does not require ATP binding.(13) Therefore the cleavage reaction of the gDNA may take place prior to N-gate closure. The yeast enzyme cannot proceed further to carry out strand passage unless ATP or an ATP analog is bound to at least one of the GHKL domains. Details of the strand passage mechanism are under active investigation but remain poorly understood.(12) After passage of the tDNA segment, the DNA gate is religated. The tDNA segment is released by dissociation of the protein:protein interface that constitutes the third gate. Finally, after ATP hydrolysis and release of ADP+Pi, the N-gate returns to an open conformation allowing release of the gDNA or capture of another tDNA segment. Interestingly, the hydrolysis of the two ATPs are sequential rather than simultaneous, and the only step that can be directly related to ATP hydrolysis is the last step in the reaction. Nonetheless, the complex coordination of the opening and closing of these three gates, and their linkage to ATP hydrolysis and gDNA cleavage, suggest extensive allosteric communication throughout the complex. However the mechanisms of these communication pathways, how they are regulated by ATP binding and hydrolysis, and how protein amino acid substitutions might affect them, remains poorly understood.

To address these allosteric mechanisms, Broeck *et al*. recently solved cryo-EM structures of the complete human Topo IIα complex bound to an ATP analogue and a gDNA segment (14). Their results showed that the human enzyme adopts an overall conformation similar to the yeast structure, including an intertwined arrangement of the subunits with the ATPase domains stacked upon the DNA-binding/cleavage domains. Notably, they observed that the ATPase domains were flexible and could adopt multiple positions relative to the DNA-binding/cleavage domains. These positions were asymmetric, with increased ATPase domain tilting in both planes relative to the remainder of the complex. This flexibility was linked to differences in the DNA-binding/cleavage domain structure, with structures in the “closed” gDNA state having a 12° additional rotation of their ATPase domains relative to those with a “pre-open” gDNA state. Furthermore, their results implicated the linker regions between the transducer and TOPRIM domains as pivotal pathways of allosteric connections. These results were in contrast to the crystal structure of the yeast Topo II structure, which showed a single conformation of the ATPase domains in a straight orientation relative to the DNA-binding/cleavage domains.

Topoisomerase mutations have played important roles in connecting structural and biochemical features of the enzyme. In particular, the genetic power of yeast based systems has led to a rich store of mutations which have been used to unveil the molecular mechanisms of TopoII enzymes. One important source of mutations have been based on screens for resistance or hypersensitivity to small molecules that target TopoII (15–17). A particularly interesting class of mutations are those that result in TopoII proteins with elevated levels of DNA cleavage in the absence of small molecule inhibitors. The first mutation in eukaryotic TopoII was found in human TopoIIα (D48N) that was found during the analysis of the action of catalytic TopoII inhibitors (18). Several other mutations which result in elevated cleavage have been recently described and have been shown to generate genome instability when expressed in yeast (19, 20). Similar mutations have been found in human TopoIIα enzymes and linked to multiple forms of cancer.(21–24)

Here, we have used molecular simulations to determine how the structure and dynamics of TopoII are regulated by nucleotide binding and hydrolysis, how they can affect the allosteric mechanisms which drive its function, and how these pathways can be disrupted by mutations leading to elevated levels of DNA cleavage. We performed a series of all-atom molecular dynamics (MD) simulations of yeast TopoII in various nucleotide-bound and mutated states. These mutants were chosen as they are located far from the active site tyrosine responsible for DNA damage, and both result in increased DNA cleavage *in vitro*. In our simulations the ATPase domains had a remarkable degree of flexibility, as suggested in the cryoEM results by Broeck *et al*., which occurs on the microsecond timescale. Furthermore, our results indicate that this flexibility and complex asymmetry are dictated by the identity of the bound nucleotide and mutations in the TopoII complex at either the amino or carboxy terminus of the catalytic core. Network analyses of these simulations identified several potential allosteric pathways via which information about the nucleotide state may be transmitted throughout the complex which involve changing the strength of communication pathways through the transducer/TOPRIM linkers. Together, our results reveal the degree to which the ATPase domains are intrinsically dynamic in TopoII and how these dynamics affect enzymatic function.

## METHODS

### Model generation

The *Saccharomyces cerevisiae* TopoII wildtype structure used in this study was obtained from the 4GFH crystal structure, which was solved in complex with the nonhydrolyzable ATP analogue AMPPNP along with a 26 base pair gDNA segmentt (11). The resolution of the 4GFH crystal structure was 4.4 Å and it was solved via molecular replacement using two higher resolution structures of the ATPase, 1PVG (1.80 Å resolution), and DNA-cleavage, 3L4J (2.48 Å resolution), domains. This resulted in a model with a good fit to the experimental crystal data, with crystalographic R_free_ and R_work_ factors of 0.277 and 0.239. In our models, the DNA was religated to form a single DNA segment that was not covalently bound to the TopoII complex, and AMPPNP was trasnformed to ATP, ADP, or deleted completely. The positions of missing residues and loops were reconstructed by using the Chimera interface to Modeller(25). Models of variants were generated by manually modifying the PDB files to introduce point mutations.

### Molecular dynamics simulations

All systems were prepared with the tleap program from the AmberTools software package (26). Systems were solvated in an orthorhombic box with a minimum distance of 10 Å to the nearest box edge and ionized with enough Na^+^ and Cl^-^ ions to neutralize the system and create an environment of ∼150 mM NaCl. The AMBER14SB, BSC0OL15, TIP3P, and Joung Cheatham parameters were used for the protein, DNA, water, and ion force fields, respectively (27–31). System sizes were ∼360,000 atoms. The ParmEd program was used for hydrogen mass repartitioning which allowed the use of a 4fs timestep(32, 33). Simulations were performed in the constant pressure/temperature ensemble, using a Monte Carlo barostat with a one atmosphere target and a Langevin thermostat set at 300 K (34, 35). Electrostatic interactions were treated with the Particle-Mesh Ewald (PME) method with a 12 Å cut-off between direct and reciprocal space (36). Systems were minimized twice for 5000 steps, first with a 10 kcal·mol^-1^·Å^-2^ harmonic restraint applied to the solute and then followed by no restraints. Initial velocities were obtained from a Maxwell-Boltzmann distribution at 5 K and heated to 300 K over 10 ps, and restraints were initially placed on heavy atoms that were gradually reduced over 1 ns. The reduction of these restraints allowed the solvent molecules to equilibrate around the solute and created a well-hydrated protein/DNA interface. Production runs were performed in quintuplicate for 1000 ns with the CUDA-enable PMEMD MD engine on a mix of local resources and those provided by the Extreme Science and Engineering Discovery Environment (XSEDE) (37, 38). The calculation of five production runs per system was done to increase reproducibility of the overall results as well as improve the reliability of error estimates throughout the manuscript.(39)

### Analysis methods

Root-mean-square-deviation (RMSD) and root-mean-square fluctuation (RMSF) analyses were performed with the CPPTRAJ package (40). RMSD was performed on all solute heavy atoms, whereas RMSF was performed for protein Cα atoms for the final 750 ns of simulations. RMSFs are reported as an average over ten value (five simulations with two subunits per simulation) with errors reported as the standard error of the mean, with a sample size of ten. dCNA network analyses were formed using R code from the Hamelberg group which utilized the Bio3D package (41–43). Molecular representations were generated with VMD and PyMol, and other figures were generated with python (44, 45).

To define the ATPase domain sampling angles of twist, tilt, and rock, the following algorithm was implemented. First, an internal coordinate system was defined based on three orthormal vectors: *X*’, *Y*’, and *Z*’. *X*’ was defined as a vector from the center of mass of the first six DNA base pair heavy atoms to the last six DNA base pair heavy atom. A central plane was defined by taking the cross-product of *X*’ with a second vector that defined the separation of the center of mass of the heavy atoms of the TOPRIM, winged-helix, and tower domains of one monomer with the corresponding atoms in the second monomer, and the normal to this plane was defined as the second axis *Y*. The third axis, *Z*’, was chosen as the cross product of the two: *Z*’ = *X*’ × *Y*’. A vector defining the orientation of the ATPase domain, *ν*, was then defined as the vector between the center of mass of the heavy atoms of the ATPase domains of one monomer with the second monomer (see Figure S3 for graphical depictions of *X*’, *Y*’, *Z*’, and *ν*). Finally, the twist was computed as the angle of *ν* with the *Y*’*Z*’ plane, tilt was computed as the angle of *ν* with the *X Z*’ plane, and rock as the angle of *ν* with the *X*’*Y*’ plane. Normalized mutual information scores between angles were computed by:

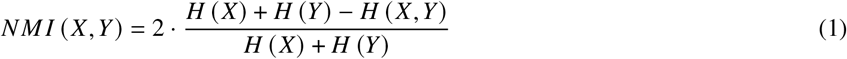

Where *H* (*X*) and *H* (*X, Y*) represented the one and two dimensional entropies of the variables *X* and *Y* .

Evolutionary conservation scores were obtained from the ConSurf database using the 4GFH PDB structure (46, 47). The ConSurf database computes evolutionary conservation scores based on a nonredundant database of homologous proteins and assigns conservation scores on a scale of one (least conserved) to nine (most conserved) using an empirical Bayesian algorithm (48).

## RESULTS

### Nucleotide States and Mutations Affect TopoII Dynamics

DNA hyper cleavage mutants perturb the breakage/reunion equilibrium during a TopoII catalytic cycle. Two hyper cleavage mutants, D48N of human TopoIIα and R1128G of yeast TopoII occur, far from the catalytic core of the enzyme, and therefore likely act by allosteric mechanisms. The budding yeast ortholog of human TopoIIα D48N is D26N. The purified protein shows elevated DNA cleavage in the absence of inhibitors (Figure S1), although its effects are less than those reported for TopoIIα D48N. Expression of yeast D26N is not lethal in rad52-yeast cells, as was observed when TopoIIα D48N is expressed in yeast, although the growth in rad52-yeast cells is markedly reduced (Figure S2). Interestingly, the yeast D26N/R1128G double mutant cannot be viably expressed in repair proficient yeast cells, suggesting the two mutations exert a synergistic effect on yeast TopoII (Figure S2).

Having established that amino acid substitutions may exert allosteric influences in yeast TopoII, we sought to examine the effects of amino acid substitutions and nucleotide bound states on TopoII structures and dynamics. To do so, we designed six simulation setups based on the crystal structure of the full length *S. Cerevisiae* system. Three of the systems had the wild type enzyme, with either two ATP, two ADP, or no nucleotides (apo state) bound. In the other three systems ATP was present in both active sites, however point mutations were introduced. These mutations were either to modify both Aspartic Acid 26s to Asparagines (D26N), both Arginine 1128s to Glycines (R1128G), or a double mutant in which both the D26N and

### ATP ADP Apo

R1128G mutations were performed (D26N/R1128G). For each system, five independent 1.0µs explicit solvent MD simulations were performed. In general the simulations were stable, with most heavy atom root-mean-square-deviations (RMSDs) ranging between 2.5 and 8.0 Å depending on the simulation run (Figure S4). There was also little change in the RMSD values after 250ns, indicating that 250ns is an appropriate equilibration time for these systems.

In these simulations the largest structural fluctuations appeared in the ATPase domains in all systems. In the ATP-bound/wildtype case, root-mean-square-fluctuations (RMSFs) of residues in the GHKL domain were the highest with fluctuations on the order of 2-4 Å (Figure 2). These RMSFs were reduced in the transducer and DNA-binding/cleavage domains, where RMSFs were typically below 2 Å, except for residues Pro562 to Lys570 which had RMSFs ranging from 2.5-4.5 Å and constitute a β-strand in the TOPRIM domains. In systems with either ADP, no nucleotide, or mutations, the RMSFs were typically lower in the ATPase domains by ∼0.5 Å, although they were roughly similar in the DNA-binding/cleavage domains, except for the previously mentioned P562-K570 β-strand (see difference of RMSFs with the ATP case, Figure S5).

**Figure 2.**
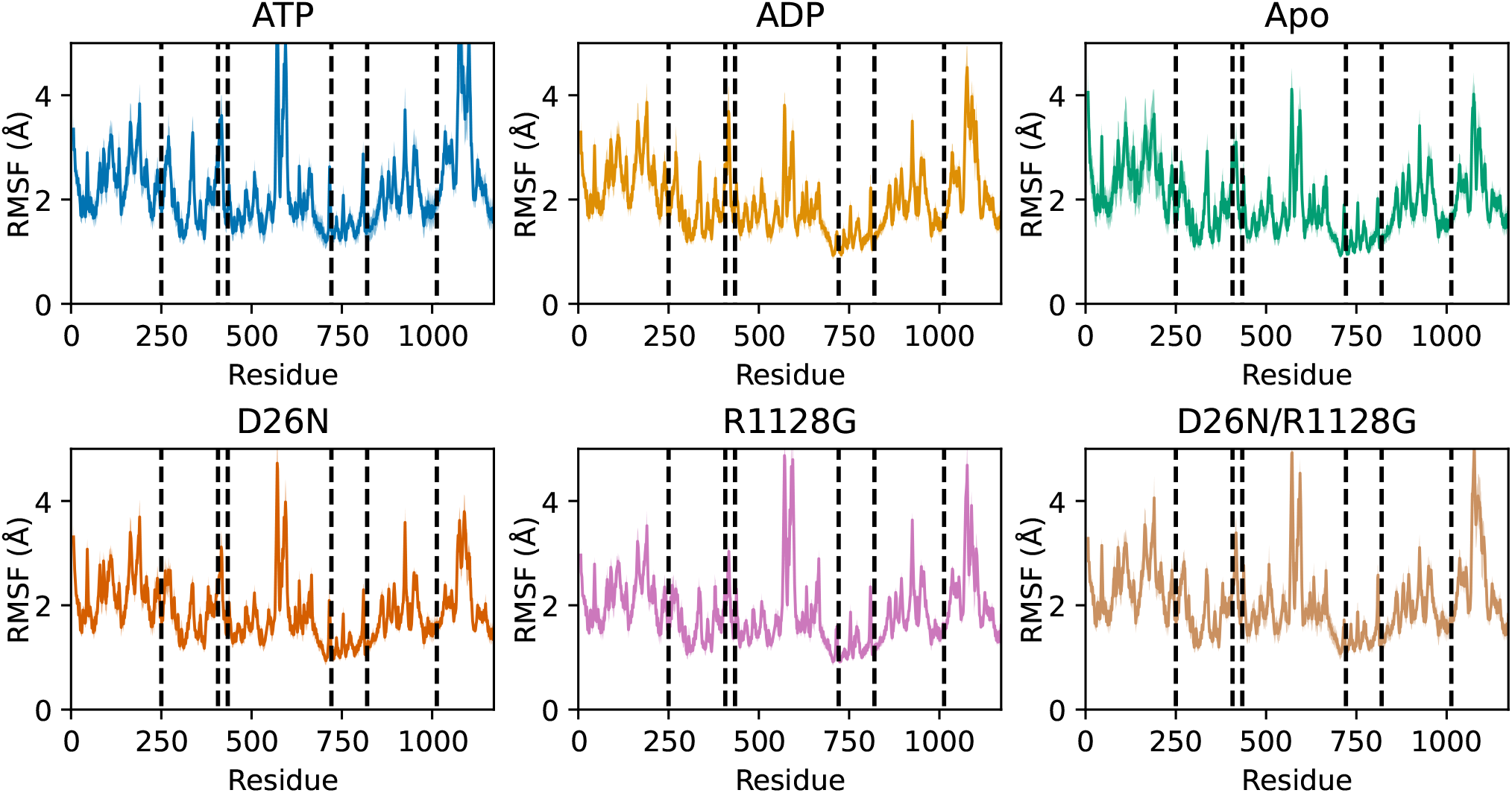
Average root-mean-square-fluctuations (RMSFs) of protein C_*α*_ atoms over the last 750 ns of each monomer of each simulation. Domains defined in Figure 1 are separated by dotted lines. Standard errors of the mean are presented in shaded colors, and protein domains are separated by dotted lines.

To understand the basis for the differences in RMSDs and RMSFs between systems, a principal component analysis (PCA) was performed on the protein C_*α*_ and DNA phosphate atoms. The results showed that the first three PCA modes captured a total of ∼90% of the observed simulation dynamics (72% for mode 1, 14% for mode 2, and 7% for mode 3), and that these motions were primarily focused on the motions of the ATPase domains relative to the DNA binding/cleavage domain (Figure 3). Although differences were noted in the sampling distributions of these PCA modes, it was difficult to use this analysis to describe the precise effects of nucleotide binding and amino acid substitutions on TopoII simulation dynamics, as the motions of each PCA mode were a complicated combination of simpler motions (such as twisting and tilting of the ATP domains). Therefore, as discussed in the next section, we developed an alternative description of the ATPase domain dynamics that allows for a more intuitive understanding of the three large-scale ATPase domain large-scale dynamics.

**Figure 3.**
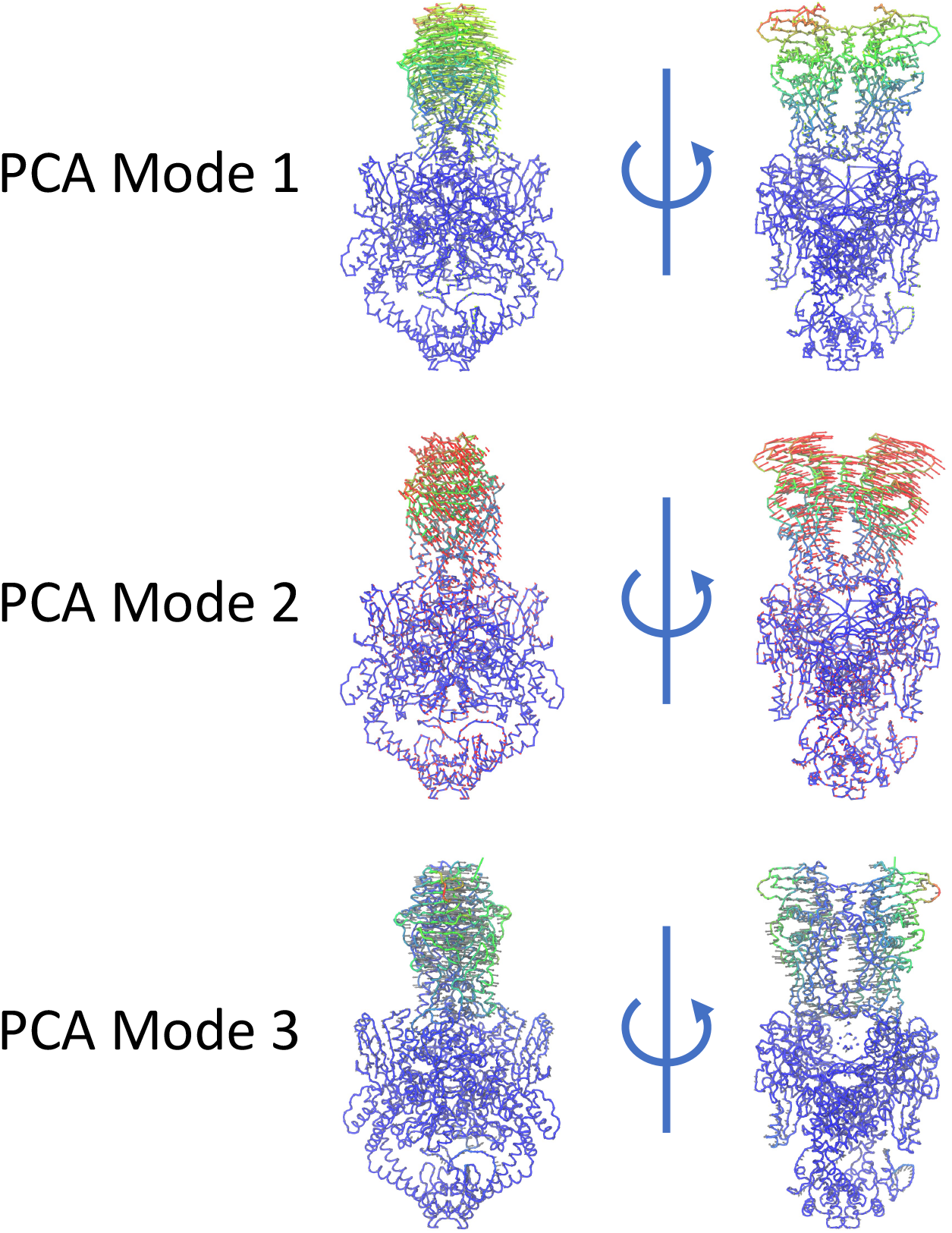
“Porcupine” plots of the first three modes from a principal component analysis (PCA) of all simulations. Protein colors represent the degree of mobility in that mode, with blue to red representing low to high motions. The arrows in each plot show the direction of motion for each residue in that mode.

### ATPase Domain Flexibility is Modulated by Nucleotide States and Allosteric Substitutions

Having found that the ATPase domains constitute the majority of TopoII simulation dynamics, we sought to decompose and quantify their motions by defining three angles: twist, tilt, and rock (see Methods for definitions, and Figure 4A for graphical depictions). The twist angle corresponds to rotation of the ATPase domains about the TopoII long axis, the tilt reflects tilting of the ATPase domains in a plane orthogonal to the gDNA, and the rock angle the tilting of the ATPase domain in a plane parallel to the gDNA (that is, orthogonal to the tilt). The initial angles for all systems had values of 76° for twist, 17° for tilt, and 0° for rock (Table 1), indicating that the ATPase domains began each simulation in a somewhat twisted and relatively upright orientation relative to the DNA-binding/cleavage domains.

**Table 1:**
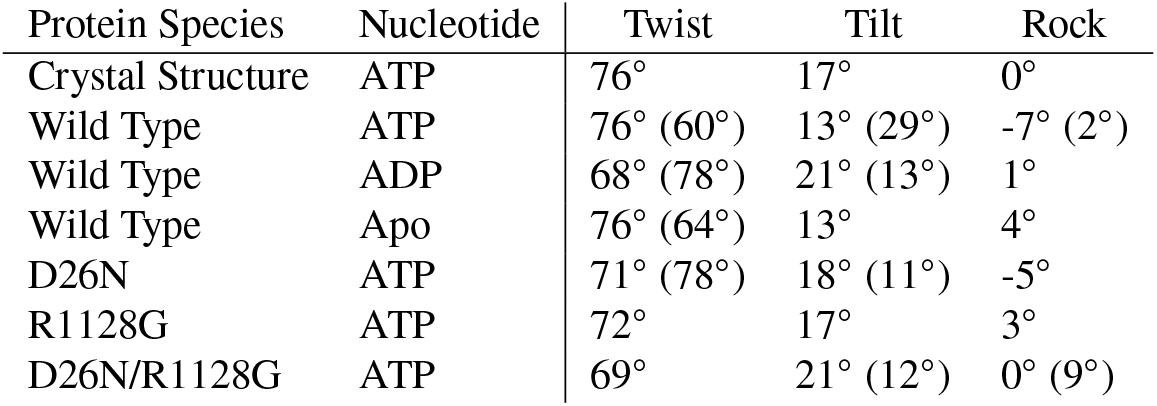
Most probable sampled angles for the twist, tilt, and rock angles of the ATPase domains sampling for each simulation setup. If multiple distinct states were sampled, the location of the second most probable value is shown in parenthesis.

| Protein Species | Nucleotide | Twist | Tilt | Rock |
| --- | --- | --- | --- | --- |
| Crystal Structure | ATP | 76° | 17° | 0° |
| Wild Type | ATP | 76° (60°) | 13° (29°) | -7° (2°) |
| Wild Type | ADP | 68° (78°) | 21° (13°) | 1° |
| Wild Type | Apo | 76° (64°) | 13° | 4° |
| D26N | ATP | 71° (78°) | 18° (11°) | -5° |
| R1128G | ATP | 72° | 17° | 3° |
| D26N/R1128G | ATP | 69° | 21° (12°) | 0° (9°) |

**Figure 4.**
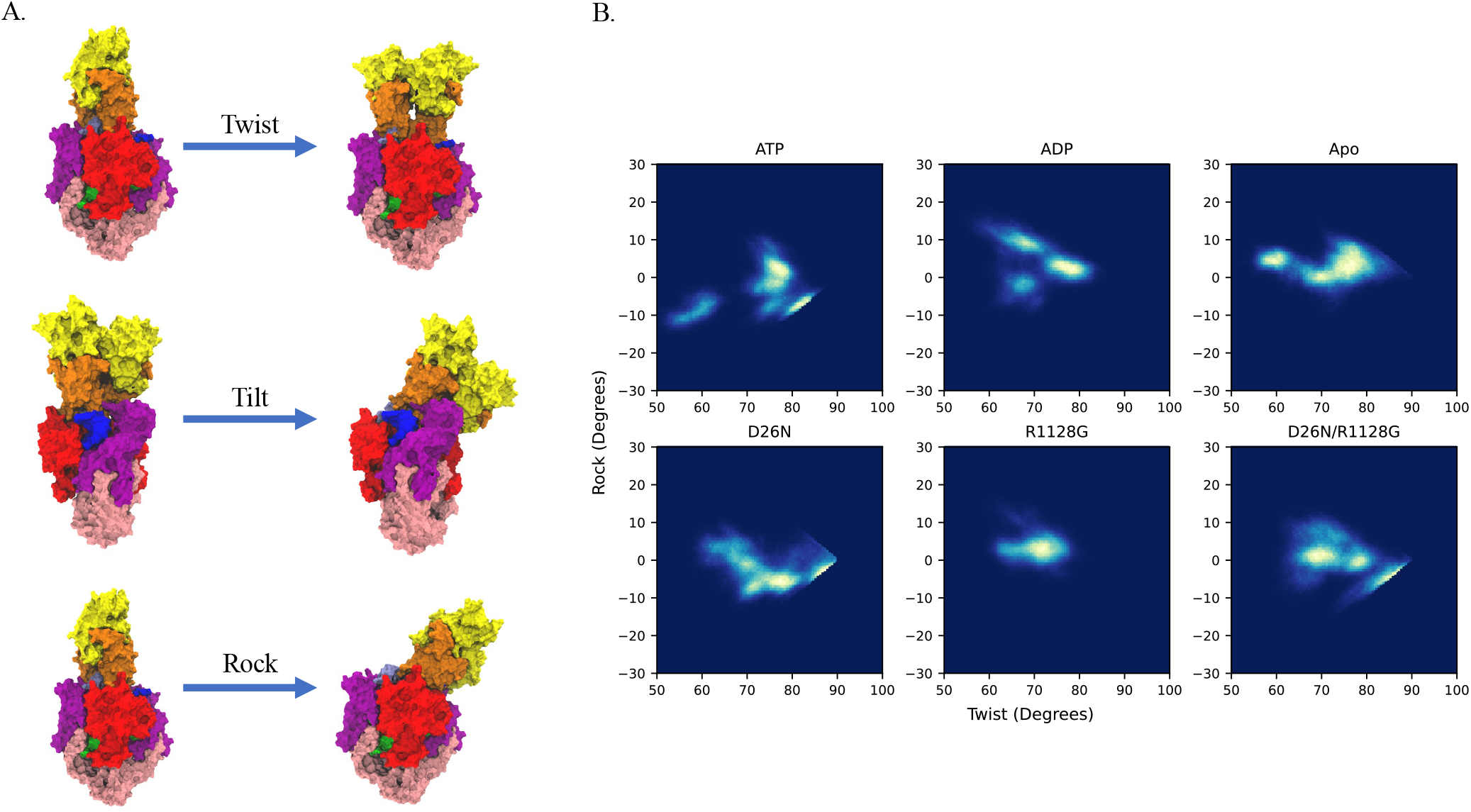
A. Depiction of the tilt, twist, and rock angles defined to describe the motions of the ATPase domains relative to the TopoII molecule (see Figure S3 for depiction of vectors used to define these angles). B. Two-dimensional histograms of the twist/rock sampling angles for each simulation setup. Results show that in the wild type/ATP case two distinct energy wells are observed, one with in an undertwisted/low rock state and another with larger twist and rock values. Nucleotide exchange or removal or amino acid substitutions alter or eliminate the presence of these states.

We observed that sampling of twist, tilt, and rock were dependent on the species of the bound nucleotides and amino acid substitutions (Table 1 and Figures S6&S7). In the ATP bound structure there was additional tilting from the crystal structure, with a local maximum in the tilt angle of 29° and rock angles around −7° and 2°, a range of motion that is consistent with the two human TopoIIα cryoEM structures observed by Broeck *et al*.(14). Although the most probable twist angle was nearly identical to the crystal structure, there was a second most probable twist value of 60°, and twist values of 49°-88° were observed, indicating that thermal fluctuations result in transient reduction of the twist angle of this domain in simulations. Upon replacement of ATP with ADP the most probable tilt angles were located around 21° and 13°, and the distribution of the twist angles had a most probable peak near the crystal structure value of 68°. Removal of the nucleotide returned the most probable twist, tilt, and rock angles to values close to the crystal structure. The amino acid substitution that had the most significant effect on sampling was R1128G, as in both the single and double D26N/R1128G substitutions the high-tilt angles were reduced relative to the ATP case to values nearly identical to the crystal structure and nucleotide free states. Both the D26N and R1128G substitutions increased the range of sampled twist values to 57°-89° and 57°-82°, which were similar to the range of 58°-90° in the double substitution.

Analysis of the two dimensional sampling space showed a coupling between these angles in the wild type/ATP bound case, and that the coupling with the rock angles was disrupted or completely abolished by nucleotide removal or amino acid substitution in our simulations. To quantify these correlations, we computed the normalized mutual information (NMI) between each pair of angles for each system (Table 2). These values show high correlations between twist and tilt in all systems, with NMI values ranging from 0.68 to 0.80. Two dimensional histograms of these angles for each system also show a nearly linear, inverse relationship between the two (Figure S8). In the wild type/ATP bound case there was a correlation between the twist/rock and the tilt/rock, with MI values of 0.21 and 0.20. These correlations were largely maintained in the wild type/ADP bound case, with values of 0.16 and 0.21, however when the nucleotide was removed or amino acid substitutions were introduced the NMI values fell such that little or no correlation was observed. The twist/rock distribution of the wild type/ATP bound case shows two energy wells: one in which the ATPase domains are undertwisted around 60° with rock values around −10°, and another with larger twist values around 80° and increased rock of −10° to 10°. Replacement of ATP with ADP or removal of the nucleotides eliminated this low-twist/low-rock state and created states with higher rock values. All mutations also eliminated sampling of this low-twist/low-rock state and created sampling distributions more similar to the ADP and nucleotide free states. Due to the high correlation between twist and tilt, the tilt/rock distribution was highly similar to the twist/rock distribution (Figure S8). In addition, comparisons of the twist, tilt, and rock two-dimensional spaces from individual simulations show similar sampling between the five simulations performed for each system (Figures S10-S12) indicating that these simulations were sufficiently converged to for this and other analyses presented here throughout the text. Overall, these results demonstrate that amino acid substitutions and nucleotide binding affect not only the range of ATPase domain motions in simulations, but also how distinct motions are coupled to one another.

**Table 2:** Normalized mutual information values between each pair of twist, tilt, and rock angles of the ATPase domains sampling for each simulation setup. Results show that in all cases the twist and tilt are highly correlated with one another. In the wild type/ATP system the rock is correlated with the twist and tilt, however this correlation is lost upon changes in the nucleotide state or protein mutations.

| Protein Species | Nucleotide | Twist/Tilt | Twist/Rock | Tilt/Rock |
| --- | --- | --- | --- | --- |
| Wild Type | ATP | 0.68 | 0.21 | 0.20 |
| Wild Type | ADP | 0.69 | 0.16 | 0.21 |
| Wild Type | Apo | 0.80 | 0.08 | 0.07 |
| D26N | ATP | 0.75 | 0.13 | 0.14 |
| R1128G | ATP | 0.74 | 0.02 | 0.03 |
| D26N/R1128G | ATP | 0.74 | 0.10 | 0.11 |

### Nucleotides and Mutations Affect TopoII Allosteric Communications

Having established that nucleotide presence and amino acid substitutions have long range effects on topoII structures and dynamics, we sought to determine the mechanisms by which those allosteric signals propagate through the system. To do this, we employed the difference contacts network analysis method (dCNA).(41, 49) dCNA is a graph-theory based approach which has the important advantage over other graph theory techniques in that it builds a consensus network across many different MD simulation ensembles. This allows for the direct measurement for how changes in the simulation setup, such as nucleotide replacement or amino acid substitutions, affect allosteric communication. Briefly, the dCNA method has four steps: 1. Contact frequencies between residues are determined across the equilibrated portions of the MD simulations for each system (here, the last 750 ns of the five MD simulations); 2. A consensus network is constructed with edge weights based upon the stability of contacts across all systems (here, the three different nucleotide states of the wildtype and the ATP bound mutants); 3. The Girvan-Newman algorithm is used to partition the consensus network into groups of residues that represent local communities of residues that are dynamically linked with one another; 4. Individual dCNA graphs are constructed by subtracting the contact probabilities of one state from another based on their common community structures.

We used the dCNA analysis to partition topoII into 20 dynamic communities. These communities typically align well with complete or partitions of the topoII subdomains (Figure 1), and in most cases are largely symmetric between the two monomers, which we denote as monomers A and B for clarity (Figure 5A). For example, communities 1 and 2 (blue and red in Figure 5) roughly correspond to the GHKL domains of subunits A and B, and 12 and 3 (purple and grey) to the transducers in these subunits. Interestingly, the exception to this was at the C-gate in which both coiled coil domains partitioned into a single community (the pink colored community 10), indicating that in the closed C-gate state the significant number of inter-subunit contacts between the coiled coil domains created a single dynamically stable community. We note that we chose 20 communities as a compromise between complexity and interpretability. When the system was coarse-grained into only 14 communities the GHKL and transducer domains of each monomer were represented by only a single community (Figure S13), suggesting a tight coupling between these divisions in the ATPase domains. In contrast, even when 26 communities were computed (the number that corresponds to the maximum modularity) the coiled-coil domains community was never split into multiple smaller communities (Figure S14), further suggesting that the residues within these units function together as a strong dynamical unit.

Changes in the contact probabilities during the ATP hydrolysis cycle demonstrate that nucleotide presence affects connections between dynamic communities throughout the complex. Replacement of ATP by ADP resulted in an increase in contacts between the two GHKL domains and a disparate effect on the transducer domain contacts (Figure 5b): the transducer in monomer B (community 3) had increased contacts with the monomer A GHKL domain (community 1) and no change with its own monomer B GHKL domain (community 2), while the monomer A transducer (community 13) had decreased contacts with both GHKL domains. In general there was an increase in contacts between communities in the top of the DNA-binding/cleavage domains, and a weakening of contacts in the bottom, including notably to the coiled-coil domains. Progression through the hydrolysis cycle of ADP being removed resulted in a significant loss of contacts between communities throughout the topoII structure, particularly in the ATPase domains and the DNA-binding/cleaving domains (Figure 5c). Completion of the hydrolysis cycle requires insertion of ATP into the nucleotide free states (Figure 5d), and generally involves a strengthening of communities within the ATPase domains, except for directly between the two GHKL domains, and a general weakening of interactions within the DNA-binding/cleavage domain to the tower domains.

**Figure 5.**
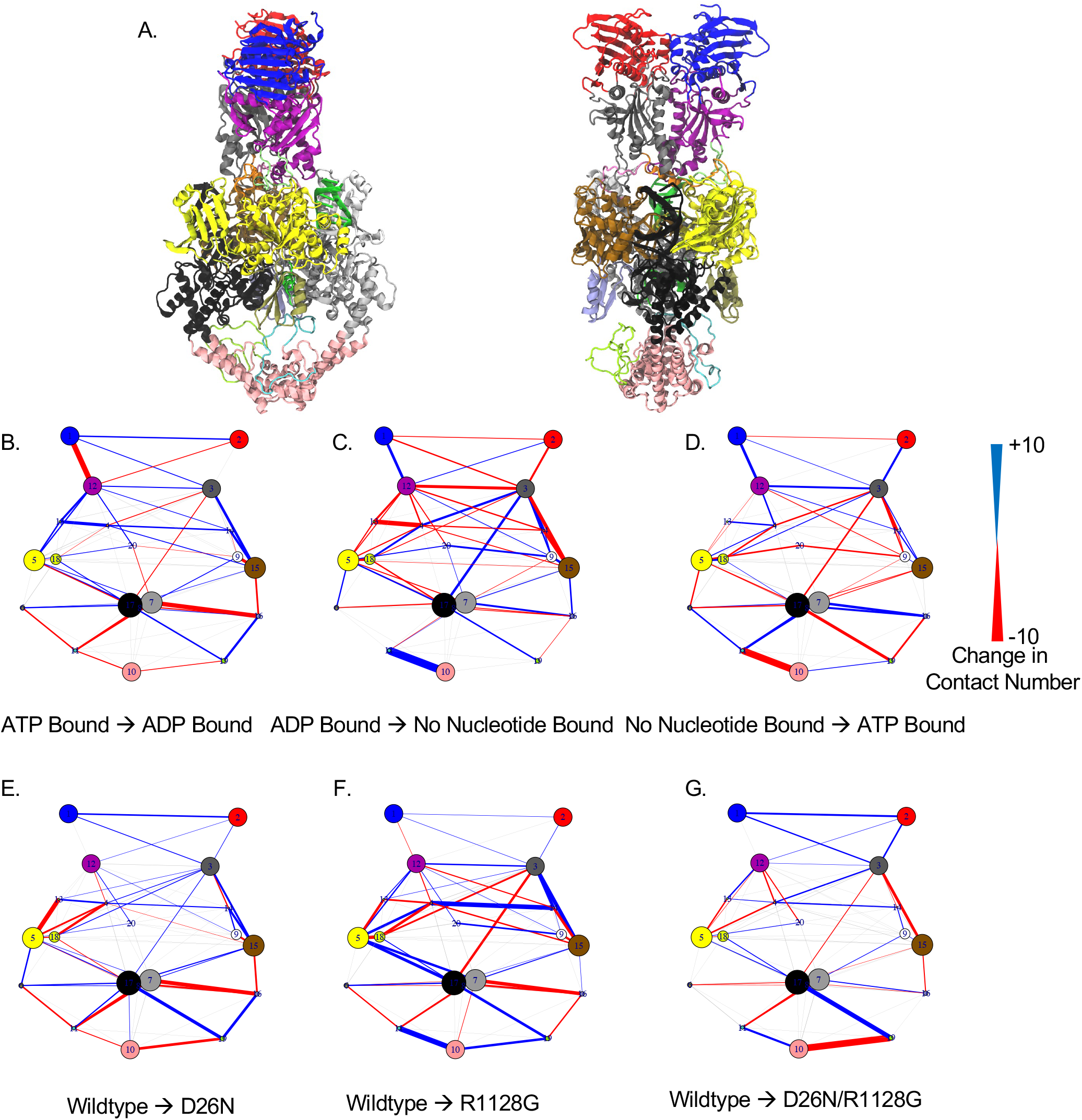
Difference contact network analysis (dCNA) results. **A**. The topoII structure was partitioned into 20 dynamic communities which roughly correspond to complete or partitions of the standard TopoII subdomain assignments (Figure 1) for the nucleotide binding cycle from ATP to ADP bound (**B**.), ADP to no nucleotide bound (**C**.), and no nucleotide to ATP bound cases (**D**.). dCNA maps were also computed for differences between the ATP bound wildtype and D26N (**E**.), R1128G (**F**.), and D26N/R1128G double mutant (**G**.) states.

Mutations appear to have long-range effects in the ATP-bound state that have similarities to the effects caused by nucleotide replacement. For example, when the D26N substitution was introduced, there was a strengthening of interactions between the GHKL domains, an increase in contacts to the tower domain in monomer B (community 17) along with a decrease to the tower domain in monomer A (community 7), and a weakening of interactions to the coiled-coil domains, all similar to the effects observed for the replacement of ATP with ADP (Figure 5e). However, unlike in the ATP to ADP case, there were increased interactions between the GHKL and transducer domains, suggesting an overall tighter ATPase domain. On the other hand, the R1128G mutation (Figure 5f) appears to create network maps more similar to what would be observed for the ATP to no nucleotide case (the opposite of Figure 5d). In that case there was a strengthening of contacts to the coiled-coil domains and a slight increase in contacts between GHKL domains. The D26N/R1128G double substitution acted as a hybrid of the two individual substitutions, with weakened interactions in the ATPase domains and a mixture of increased and decreased interactions to the coiled-coil and tower domains (Figure 5g).

### Pathway Analysis Suggest the Molecular Mechanisms of Allosteric Communication

To probe the mechanisms that underlie these long-range allosteric effects, we performed a series of pathway analyses based on the dCNA networks (42). In these analyses, the shortest 500 pathways between two residues were computed, with the path length between two residues defined as the change in the number of contacts upon a perturbation, such as nucleotide exchange or amino acid substitution. The path length is inversely correlated to the efficacy of communication along that pathway, with shorter lengths corresponding to stronger communication. We computed pathways between each of the Glu66 residues with each of the catalytically critical Tyr782 residues in the DNA-gate and the Thr1125 residues in the C-gate, for a total of eight pathway analyses per system. Glu66 was chosen as it makes direct contacts with the γ-phosphate of ATP and acts as a general base in TopoII and gyrase enzymes,(50–52), while Tyr782 cleaves and binds to the G-DNA strand.(11) Although Tyr1125 is not critically crucial for TopoII function, it is located at the end of an α-helix at the base of the coiled-coil domain and forms direct contacts with the opposing monomer, and was therefore chosen as a representative location of the C-gate.

Visual inspection of the results between each Glu66 and Tyr782 showed similar pathways for each nucleotide exchange or amino acid substitution. For example, communication between each Glu66 and the Tyr782 of the same subunit initially proceeds along a well-defined path of Glu66*→*Asp225*→*Tyr254*→*Ala309 (Figure 6b&d). However, upon reaching the transducer domains the suboptimal pathways diverged and there were multiple communication pathways through the interface between the ATPase and DNA-binding domains. A funneling of these pathways occurs as the residues enter the DNA binding domain, with some pathways passing through the gDNA strand. Pathways between each Glu66 and the Tyr782 of the other subunit were also visually similar and followed the same initial pathway that consisted of Glu66 through Ala309 (Figure 6c&e). However, in both cases they diverged into two major sets of pathways before rejoining near the opposing Tyr782.

**Figure 6.**
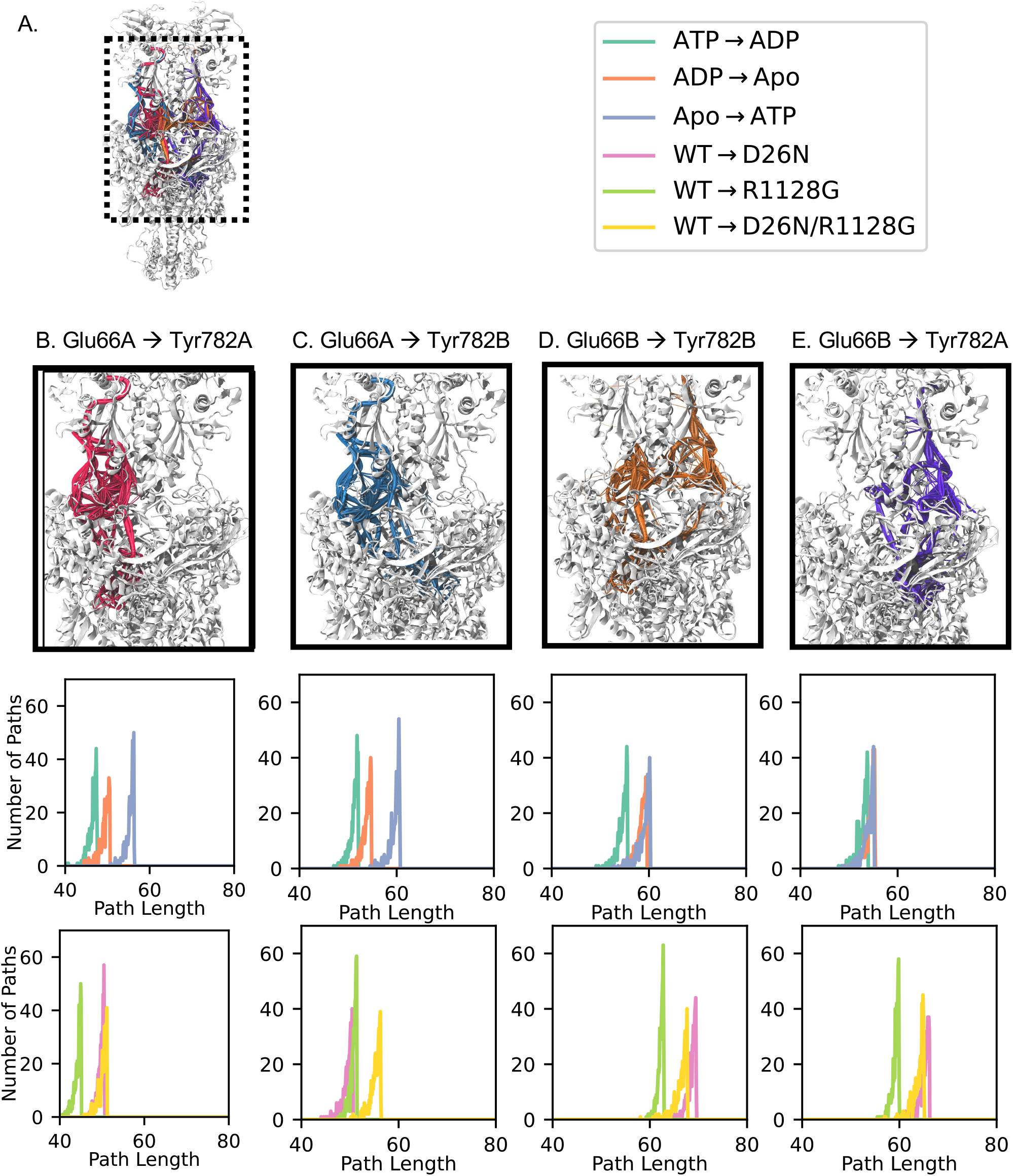
Residue suboptimal pathway analysis for networks beginning at each Glu66 residue and terminating at each Tyr782. (a) Global view of each set of pathways, along with color definitions for pathway length histograms. Pathways for (b) Glu66A→Tyr782A, (c) Glu66A→Tyr782B, (d) Glu66B→Tyr782B, and (e) Glu66B→Tyr782A, along with their respective lengths.

Although the pathways appeared similar between systems, there were significant differences in their lengths, which is reflective of the strength of these communication networks (Figure 6). For each set of pathways in the wildtype systems, the ATP*→*ADP transition had the shortest pathways, indicating that replacement of ATP with ADP is easily detected throughout the TopoII complex. Surprisingly, the Apo*→*ATP system had some of the longest pathways despite the fact that this represents a significant physical perturbation. This may reflect the fact that the closed N-gate/Apo state is somewhat artificial, and that the natural state of the Apo system is a more open system not simulated here. In addition, we note an asymmetry in the path lengths. Pathways between Glu66A and Tyr782A were significantly shorter than between Glu66B and Tyr782B, indicating that communication is stronger in one subunit than the other. Mutations appeared to exacerbate this effect, with pathways beginning at Glu66A being roughly equal in length to those in the wildtype (Figure 6b&c), whereas pathways that began at Glu66B had lengths that were significantly longer than their wildtype counterparts.

Analyses of pathways terminating at the two Thr1125 residues suggests similar overall conclusions (Figure S15). In each case, pathways begin through the same well defined route of Glu66 through Ala309, however they then diverge at the transducer domains and also throughout the coiled-coil domains. As with the wildtype systems there is an asymmetry in the path lengths, with paths beginning at Glu66A being shorter than those to that begin at Glu66B. This asymmetry increased in mutations, which created much longer path lengths in the second monomer.

One downside of pathway analysis is that it requires the definitions of endpoints. Although in some cases these endpoints may be obvious for a particular mechanism, such as the choice of Glu66 and Thr782 for examining the connection of nucleotide binding to DNA cleavage, it can miss pathways between other residue pairs which may be important for other enzymatic mechanisms. To determine which residues are most important for global communication, we computed the betweenness centrality scores for each dCNA network. In betweenness centrality analysis, the shortest path between each pair of residues are computed, and residues are ranked by how many of these shortest paths pass through each of them. This results in a scoring of the importance of each residue to the global communication networks in the system.

Betweenness centrality analysis of the dCNA networks imply that the transducer and DNA binding regions are important communication hubs in TopoII. In particular, in the network formed by the replacement of ATP with ADP, multiple residues along the linker helix have high betweenness centrality scores (Figure 7a). In addition, residues around the DNA and DNA basepairs themselves had particularly high betweenness centrality scores, indicating that the gDNA strands actively participate in allosteric communication. There was some variations in betweenness centrality scores between the different dCNA networks.

**Figure 7.**
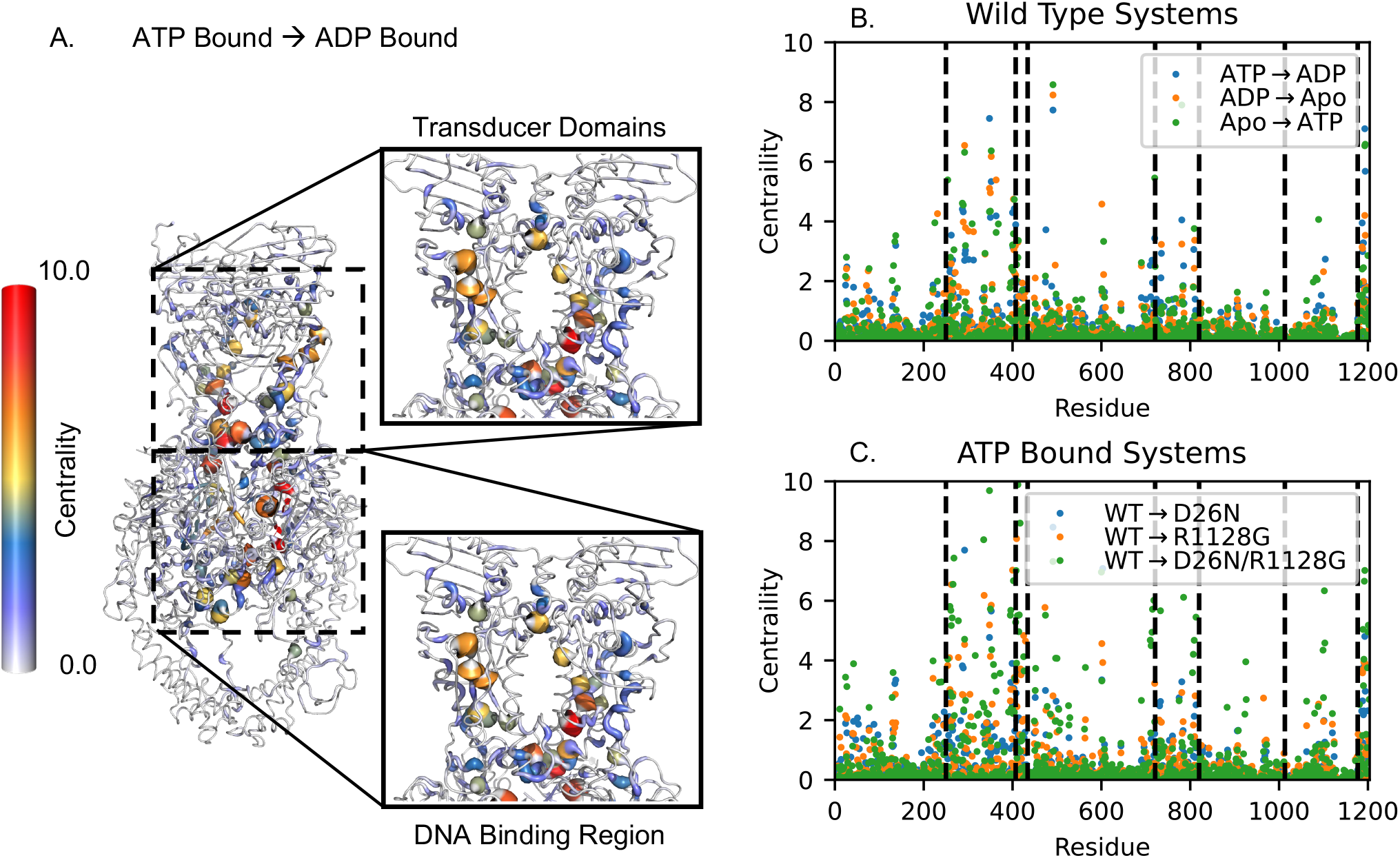
Betweenness centrality scores for networks generated through a dCNA analysis. (A.) Residues with the biggest influence on dynamics networks that are modified in the ATP→ transition are primarily located in the transducer domain and the DNA binding region. Structures representing all betweenness centrality scores are in Figure S16. (B.) Betweenness centrality scores for nucleotide modifications in the wildtype systems show the largest changes in the transducer domains, as well as the DNA and DNA binding regions. (C.) Betweenness centrality scores for mutations in the ATP bound state also show high scores in the transducer and DNA binding regions.

For example, betweenness centrality scores in the wildtype enzyme tended to be higher in the ATP*→*ADP network than in the ADP*→*Apo or Apo*→*ATP networks, further indicating that allosteric networks are particularly sensitive to the loss of the γ-phosphate (Figures 7b, S16, S17). Furthermore, mutations appear to disturb the networks in the ATP bound state to varying degrees. The R1128G mutation had the lowest effect on betweenness centrality scores in the transducer domain and DNA binding regions, however both the D26N and the D26N/R1128G had effects that were closer in scale to those observed in the nucleotide exchange systems, with the double mutant having some of the largest betweenness centrality scores with values above 8.0 (Figures 7c, S16, S17).

To determine if residues with high betweenness centrality scores were conserved in evolution, evolutionary conservation scores for TopoII were extracted from the ConSurf database, which ranks residues on a one (low conservation) to nine (high conservation) scale (46, 47). Comparisons were made between the conservation and betweenness centrality scores by extracting the upper 10% and lower 10% of betweenness centrality scores from either the ATP to ADP dCNA network, or the per-residue maximum score from all of the wild-type networks (Figure 8). In both cases, highly conserved residues are highly populated within the upper 10% of betweenness centrality scores while the least conserved residues are observed with increased frequency in the lower 10% of betweenness centrality scores. This indicates that there is some correlation between the betweenness centrality and the evolutionary conservation scores, which are, by design, measurements of different residue properties. Residues with high evolutionary and betweenness centrality scores are located mainly in the TOPRIM and WHD within the DNA-gate and transducer domain within the N-gate, indicating that these residues play a key role in both dynamic networks as well as the classic structure/function relationship (Figure S18 and Table S1). However, there are also some residues with high betweenness centrality and low evolutionary scores, or visa versa. For example, residues in the transducer linker region have high betweenness centrality scores and lower evolutionary scores, and have also been implicated in TopoIIα allosteric networks in the work of Broeck *et al*.(14). This suggests that this region of TopoII is important for dynamic communication, especially as it likely acts as a “funnel” for information transfer from the ATP-gate to the DNA-gate, however individual residues in this area may be able to tolerate multiple mutations that may not significantly alter the overall structure and dynamics of the complex. Similarly there are residues throughout the complex with high conservation but low betweenness centrality scores. These residues are likely of importance for structural stability of the complex, but may not play a significant role in the transmission of allosteric information.

**Figure 8.**
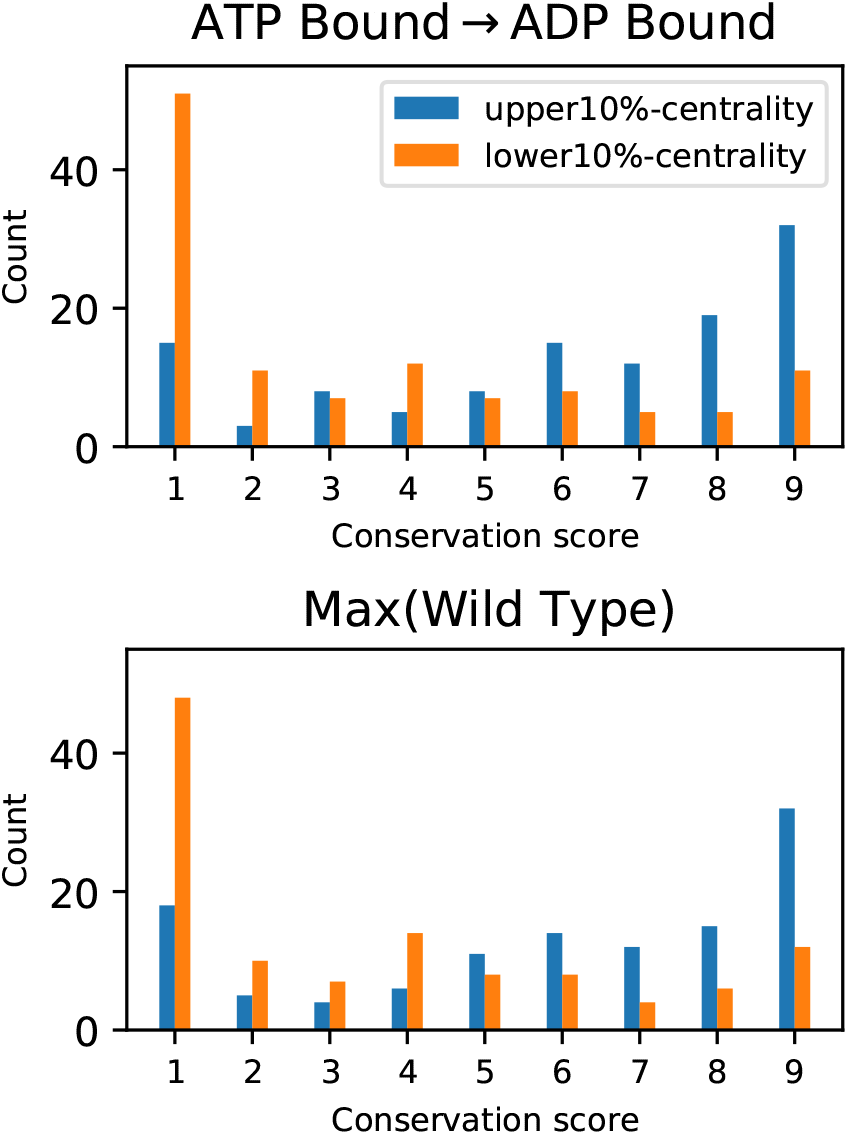
Conservation scores for the upper and lower 10 percent of residues when the betweenness centrality scores ordered in descending order. Top: Betweenness centrality scores from ATP→ADP system and Bottom: Maximum betweenness centrality scores out of three WT systems.

## DISCUSSION

The simulations presented here reveal intricate details of how *S. cerevisiae* TopoII’s microsecond timescale dynamics may be coordinated by nucleotide binding, and how they can be disrupted through amino acid substitutions. It has previously been shown in cryoEM experiments on the human TopoIIα enzyme that the ATPase domains can adopt multiple conformations in solution(14). Here, we analyzed a series of simulations on yeast TopoII systems of different nucleotide bound states, as well as those with amino acid substitutions, and indeed our results show that these domains are highly flexible in the yeast enzyme, suggesting that ATPase domain flexibility may be ubiquitous in TopoII enzymes. RMSF results show that the largest structural fluctuations tended to be present in the ATPase domains with increased motions in the ATP bound state. These dynamics involve a combination of motions, which were decomposed into twisting and tilting motions of the domains relative to the DNA-binding/cleavage domains. Nucleotide binding and amino acid substitutions resulted in systems adopting specific states, notably in the Rock/Twist space. They also resulted in altered allosteric networks throughout the complex, with the effects on the interactions of communities of residues determined via dCNA analyses and specific pathways delineated by pathway analyses. It was noted that the change from ATP to ADP in the nucleotide binding site was felt the most strongly throughout the complex, as described by the shorter path lengths between key TopoII enzymes in the dCNA pathway analyses. Using betweenness centrality analysis on these networks, we were able to identify specific residues that appear to contribute to the communication pathways in the TopoII complex, many of which are located in the Transducer domains and are evolutionarily conserved. Given that our simulations were performed on the yeast enzyme, the observation of ATPase dynamics in line with those observed in human TopoIIα suggests that ATPase dynamics are an intrinsic property of TopoII enzymes. These dynamics likely play a central role in the ATPase domains’ mechanisms of regulating the DNA strand passage mechanism, as they provided a process for coordinating the physical processes of strand capture and passage with the ATP hydrolysis.(53)

Although these simulations represent only one stage of the complete TopoII catalytic cycle, they provide a baseline for understanding how dynamical properties may affect other stages of the strand-passage mechanism. For example, when a tDNA molecule is captured by TopoII it must pass through both the DNA- and C-gates. To pass through the DNA-gate, the ATPase domains would likely need to rotate to align the tDNA to be perpendicular to the gDNA. The nucleotide dependent twisting of the ATPase domains may help to control that twisting as a means of funneling the tDNA through the gDNA. Furthermore, our network analysis showed that the coiled-coil domains of both subunits form a single, strong dynamic unit. On its face this would be counter to the idea of an opening of the C-gate which would require the breaking of inter-subunit interactions. However, the presence of tDNA below the DNA-gate would likely alter these contacts and may serve as a lever to help open the C-gate. Therefore, it is possible that the presence of the tDNA itself may be a key determinant in the opening of this gate.

Given the complexity of the TopoII cycle, and the vast range of dynamical motions that regulate it, it is notable that mutations at distant ends of the complex can have long-range effects on the catalytic sites. Here, we examined two mutations, both of which are distant from the gDNA binding region. One of these (D26N) is located in the ATPase domains, while the other (R1128G) is near the N-terminal of the coiled-coil domains. We also studied a third mutant, which is the double-mutant of D26N and R1128G. In all three cases, there was a dramatic effect on the dynamics and allosteric pathways of the complex. It is striking that both D26N and R1128G (as well as the D26N/R1128G double mutant) induce alterations in the ATPase domain dynamics that at least superficially resemble alterations that occur in the enzyme as the nature of the bound nucleotide state is changed. Since both D26N and R1128G are far from the active site or the TOPRIM domain, these mutations are likely to act via an allosteric mechanism, rather than directly participating in enzyme mediated cleavage reactions directly. In most cases, the allosteric pathways between the nucleotide-binding region and the DNA- and C-gates were weakened by mutations, despite these mutations not occurring within the identified pathways. The mechanisms of these effects are varied. In the case of D26N, mutations serve to increase contacts between the GHKL and Transducer domains, modestly decrease ATPase domain fluctuations, and increase the twist and tilt of the ATPase domains. In the case of R1128G an increase in contacts within the coiled-coil domains was noted, along with similar effects on ATPase domain fluctuations.

The simulations presented here lack the disordered C-terminal domains given their size and disordered nature. The effects of these domains on the catalytic cycle is not entirely clear. On the whole, these domains have nuclear localization signals and are subjected to a plethora of post-translational modifications *in vivo* (54–56). It was shown by Broeck *et al*. that human TopoIIα enzymes lacking the complete C-terminal domain had similar relaxation activity to the wild type enzyme, although if only a small portion of the domains were preserved there was a significant decrease in relaxation activity. This suggests that in the human enzyme there may be a complex interplay between portions of the C-terminal domains with the structured portions of TopoIIα. Given that the C-terminal domains are poorly conserved between human and yeast enzymes and the lack of structural information about these domains in yeast, we can not predict based on our current simulations how these regions would affect TopoII’s overall mechanism. Based on the location of the interface between the coiled-coil and C-terminal domains and the location of the linker region in the human TopoIIα enzyme it is tempting to speculate that these domains may make contacts with the G-strand DNA, especially if the G-strand is significantly extended beyond the relatively short segment used here. Such contacts may influence the strengths of some of the allosteric pathways observed here, notably those that include portions of the G-strand DNA. However, further experimental and computational work is required to discern these effects.

This study significantly extends recent computational work on TopoII system. In a recent simulation study by Ogrizek *et al*. on the isolated human Topo IIα ATPase domains, it was noted that the transducer domains have significant flexibility relative to the GHKL domains on the hundred nanosecond timescale. (52) This is in line with what we observed in the full complex, where this relative motion between the GHKL and transducer domains leads to large-scale motions of the ATPase domains relative to the remainder of the protein, as the base of the transducer act as a hinge for the full N-gate. Chen *et al*. presented complimentary simulations of the human TopoIIβ DNA gate without the ATPase domains in which they induced tDNA strand passage with steered MD simulations.(12) In those simulations it was shown that strand passage may occur through a “rocker-switch” mechanism. However, these studies could not determine how this process may be regulated by ATP hydrolysis due to the truncated TopoII structure lacking an N-gate region. Here, we show that hydrolysis is linked to DNA-gate dynamics through a series of allosteric networks that actively utilize the bound gDNA and which are sensitive to the identity of the bound nucleotide and the amino acid composition throughout the complex. In a simulation complementary to our own, Bandak *et al*. performed a community structure analysis on a 500ns simulation of the full-length human TopoIIβ enzyme bound to ATP.(57) Their results showed a similar partitioning of the TopoII structure into communities as ours, indicating that the conclusion that TopoII enzymes function as dynamic hubs is robust to the species and type of TopoII enzyme under consideration. In addition, they showed that hyper cleavage mutations preferentially localize to nodes between communities, which would be expected for sites that may exert allosteric effects. Our present analysis is in accord with this view. It should be noted that hyper cleavage mutants could have a more direct effect on the cleavage/religation reaction, and the recently described 720 mutation may be an example of a hyper cleavage mutant that may exert a more direct effect, given its proximity to the catalytic active site tyrosine.(20) Nonetheless, our work supports the notion suggested by Bandak et al. that hyper cleavage mutations are most likely to exert their effects by allosteric mechanisms. Future studies linking the capture, passage, and release of both the gDNA and tDNA are needed to fully determine the molecular details of each step of the TopoII catalytic cycle.

## AUTHOR CONTRIBUTION

J.L.N. and J.W. conceived the project; S.E., N.L.K., and J.W. carried out the molecular dynamics simulations and analyses; K.C.N. and J.L.N. performed the biochemical analyses and topoisomerase biochemical analyses; all authors performed comprehensive analyses; S.E., N.L.K., J.L.N, and J.W. wrote the manuscript; all authors have agreed to the final content

## ACKNOWLEDGMENTS

We thank members of the Wereszczynski group for valuable conversations concerning the simulations and analyses presented here along with Drs. Jose Villegas and Matthew Schellenberg for helpful comments on the manuscript. Figure 1c was created with BioRender.com. Work in the Wereszczynski group was funded by the National Institutes of Health [R35GM119647]. This work used Expanse system at the San Diego Supercomputer Center through allocation MCB140081 from the Advanced Cyberinfrastructure Coordination Ecosystem: Services & Support (ACCESS) program, which is supported by National Science Foundation grants #2138259, #2138286, #2138307, #2137603, and #2138296.

## DATA AVAILABILITY

MD trajectories generated in this study are available on Zenodo at: https://doi.org/10.5281/zenodo.8208867

