## Supporting Information for "Modeling Allosteric Mechanisms of Eukaryotic Type II Topoisomerases"

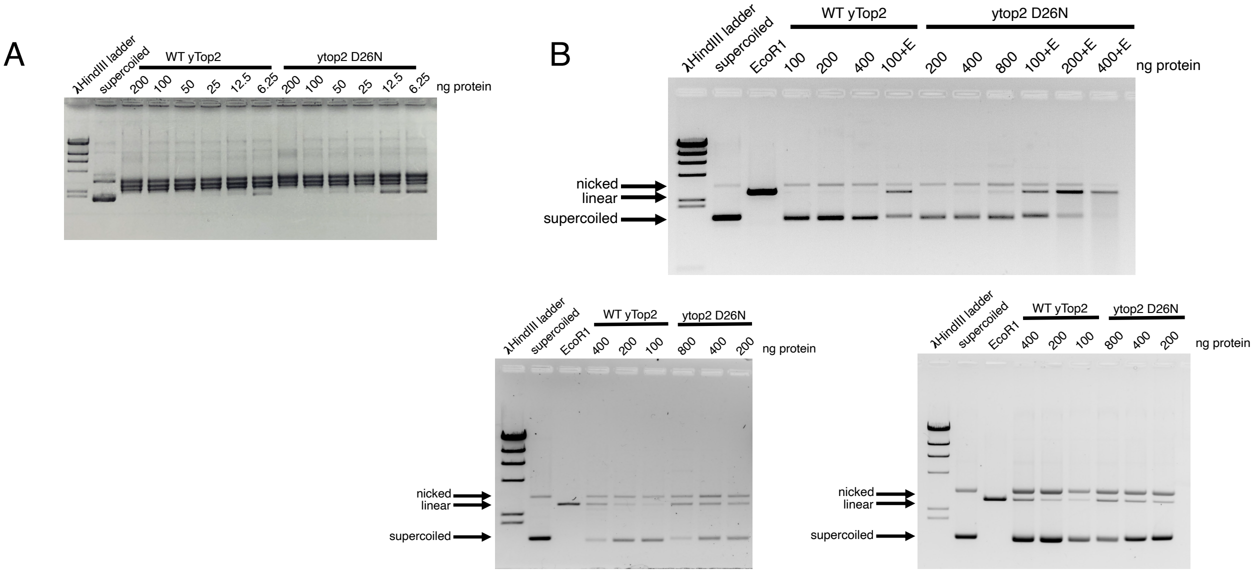


Figure S1: Enzymatic activity and DNA cleavage by yeast Top2-D26N. Wild type yeast TopoII and yeast TopoII-D26N were purified as previously described (1). (A.) Relaxation of supercoiled pUC18. Relaxation assays were carried out as previously described(2). Note that the proteins were purified from yeast cells deleted for TOP1, so there is not interference by contaminating topoisomerase 1 activity. We conservatively estimated that TopoII-D26N was at least two fold less active than the wild-type enzyme. (B.) DNA cleavage activity of wild- type yeast TopoII and yeast TopoII-D26N were carried out using standard protocols(2). The amounts of TopoII-D26N protein were chosen to reflect its lower activity. TopoII-D26N protein showed elevated levels of cleavage and elevated levels of cleavage in the presence of etoposide. The results of three independent assays are shown.

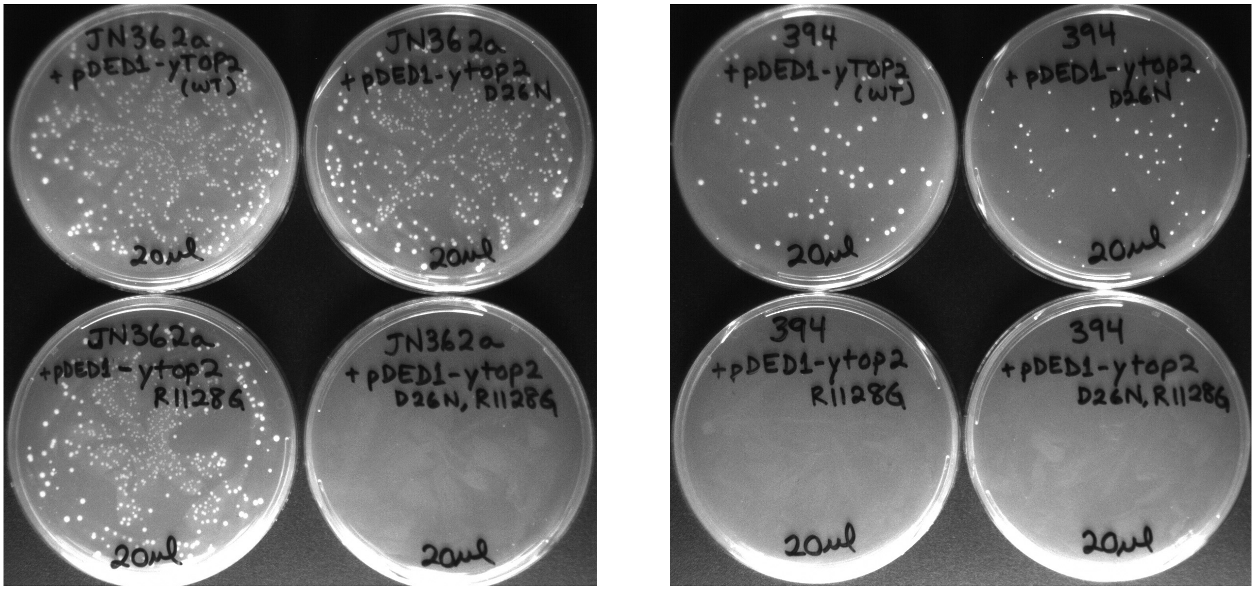


Figure S2: Growth of RAD52+ and rad52- yeast cells upon transfection with TopoII hyper cleavage alleles. RAD52+ (JN362a) and rad52- (JN394) strains have been previously described (3). Cells were transformed with plasmid pDEDyTop2 wild type, pDEDyTop2-D26N, pDEDyTop2 R1128G, or pDEDyTop2 D26N, R1128G as indicated on the labelled plates. All plasmids except pDEDyTop2 D26N, R1128G efficiently gave rise to colonies in RAD52+ cells. Interestingly TopoII D26N, R1128G mutants could not be expressed in wild type cells under our conditions. pDEDyTop2-D26N mutants could be transfected into rad52- cells but the colony number and size are reduced compared to wild type TopoII. pDEDyTop2 R1128G could not be transfected into rad52- cells, in agreement with our previously published results(4). Since TopoII D26N, R1128G mutants could not be expressed in wild type cells, it is unsurprising that they could not be expressed in rad52- cells. The double mutant protein has not yet been purified, but we suggest that it likely has further elevated DNA cleavage. The inability to express hyper cleavage mutants in wild type yeast cells has been previously seen with human TopoIIα-K743N(5) .

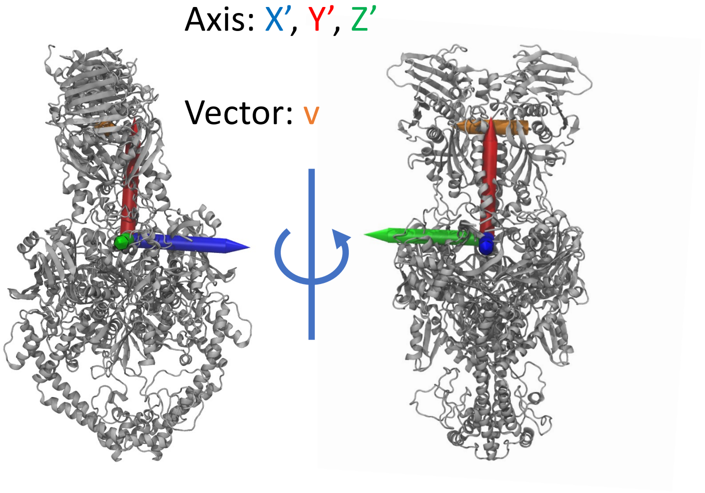


Figure S3: Overlay of angles $X'$, $Y'$, $Z'$, and $v$ used to define the tilt, twist, and rock angles.

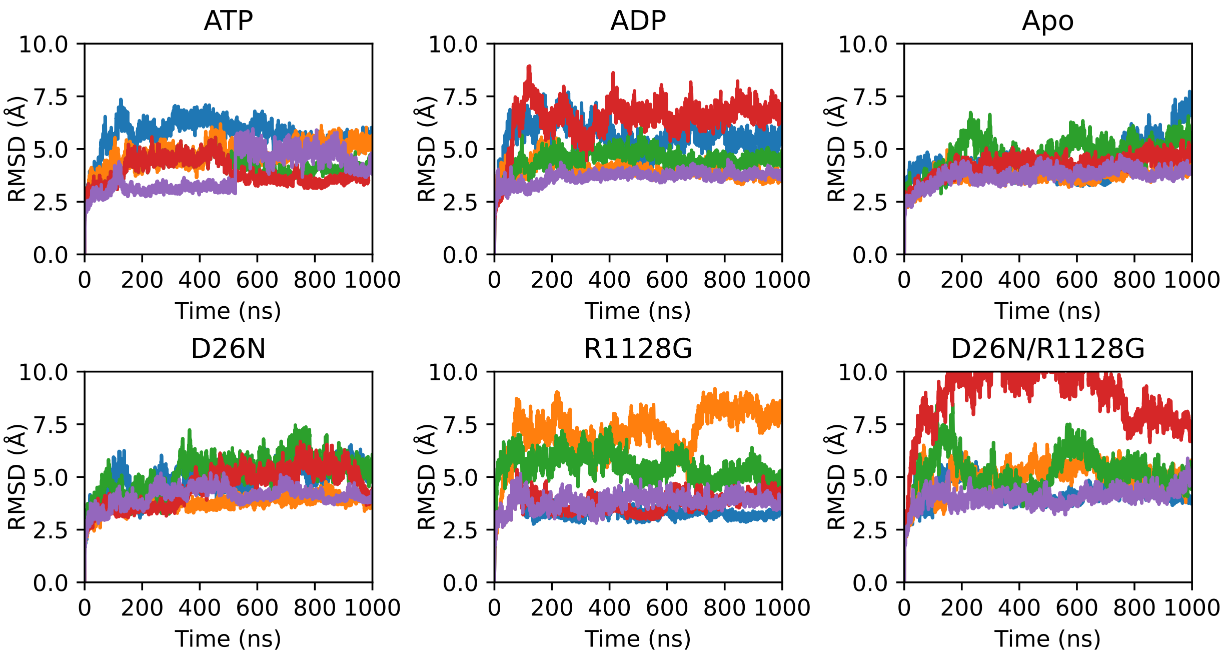


Figure S4: Root-mean-square-deviations of the heavy atoms for each simulation. In each plot the colors represent separate MD runs.

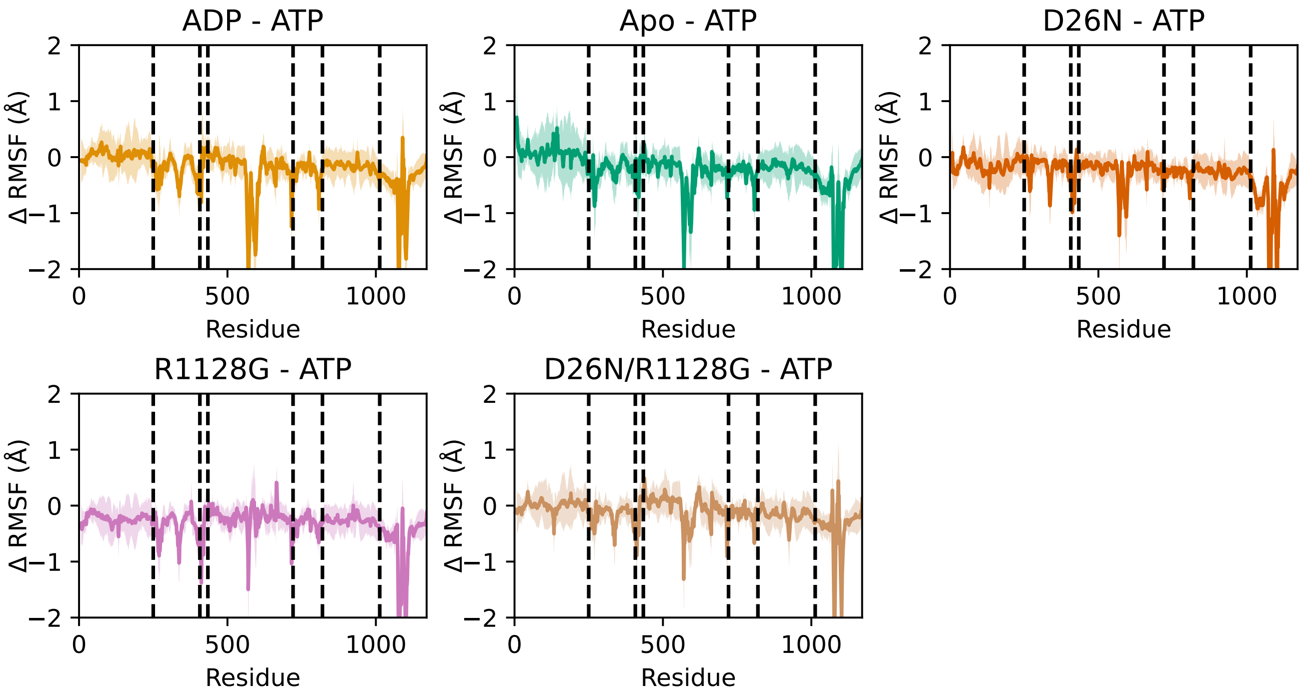


Figure S5: Difference of protein $C_\alpha$ RMSFs in Figure 2 to the ATP case. Standard errors of the mean are presented in shaded colors, and protein domains are separated by dotted lines.

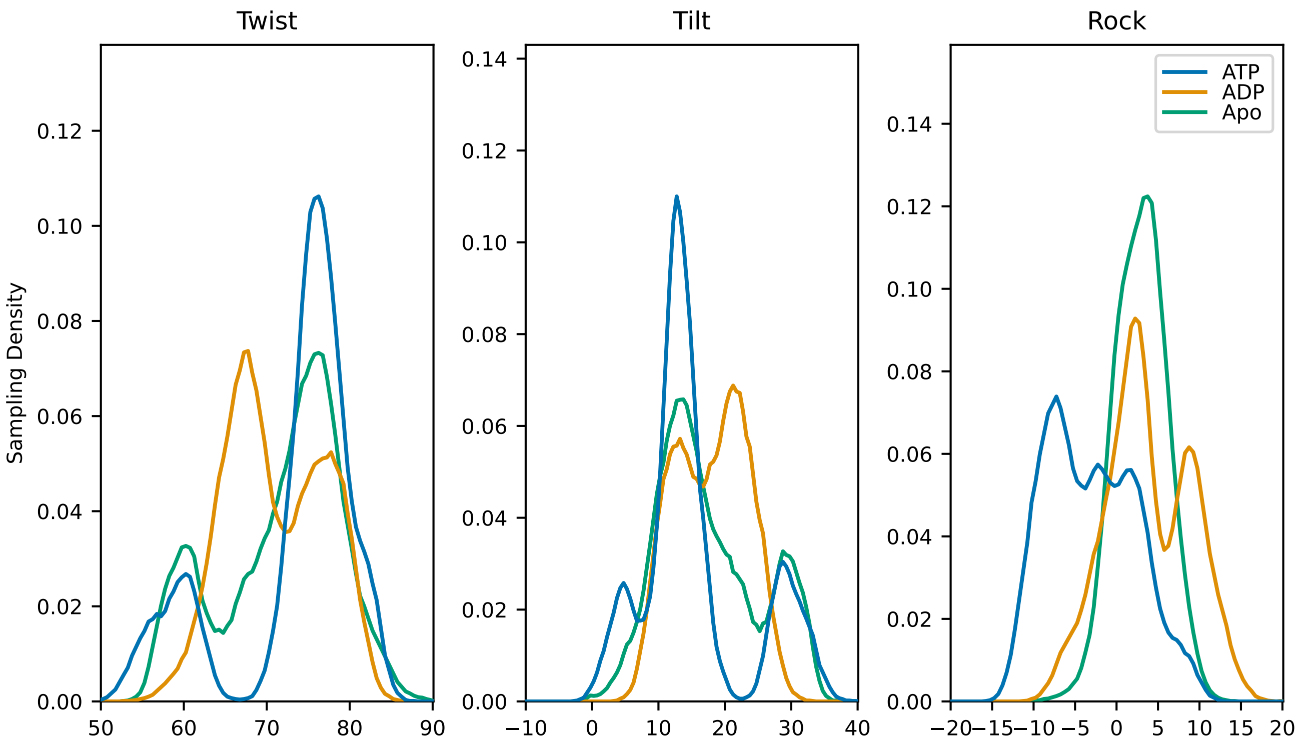


Figure S6: One dimensional distributions of twist, tilt, and rock sampling for the wild type systems with different bound nucleotides.

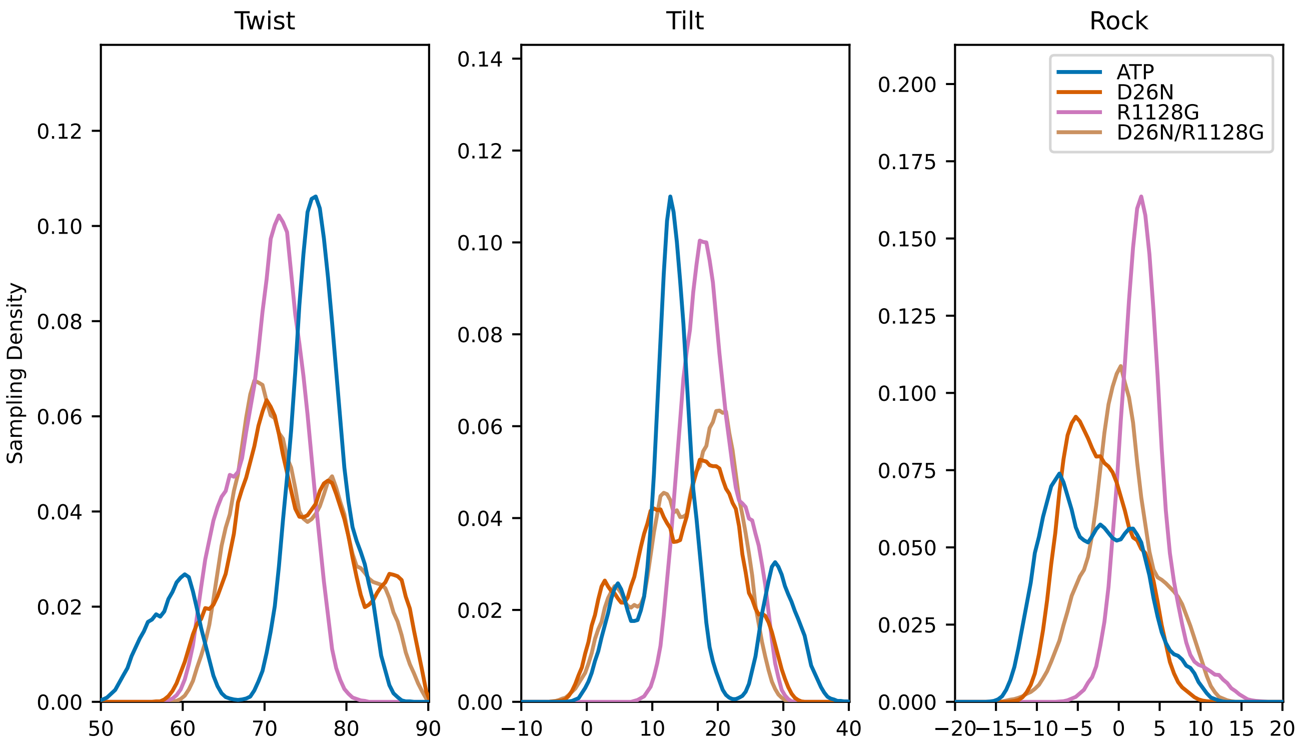


Figure S7: One dimensional distributions of twist, tilt, and rock sampling for the wild type and mutation systems with ATP bound.

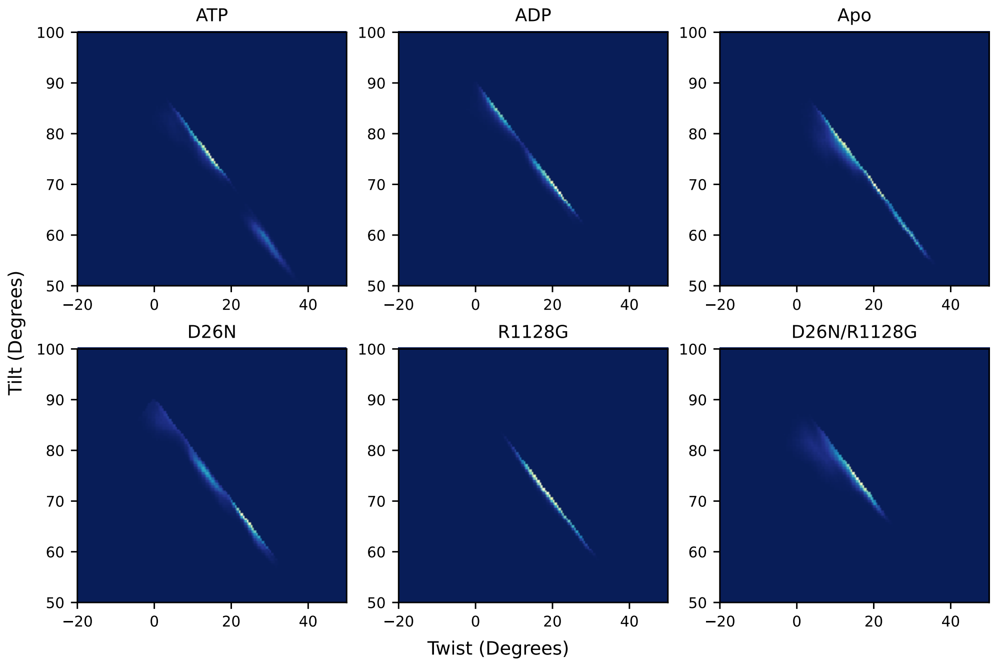


Figure S8: Two-dimensional histograms of the twist/tilt sampling angles for each simulation setup.

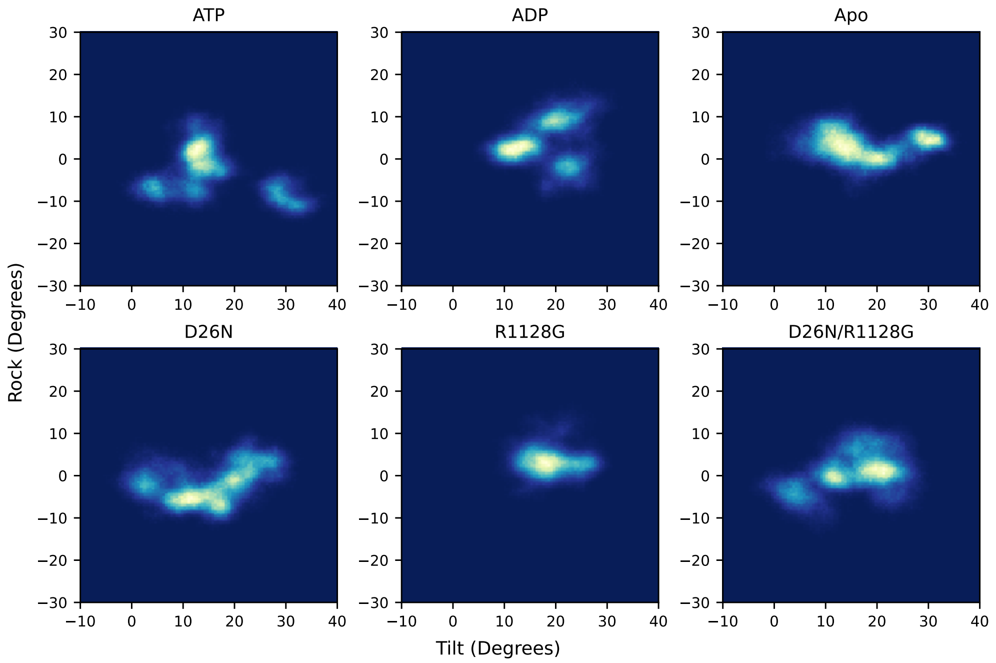


Figure S9: Two-dimensional histograms of the tilt/rock sampling angles for each simulation setup.

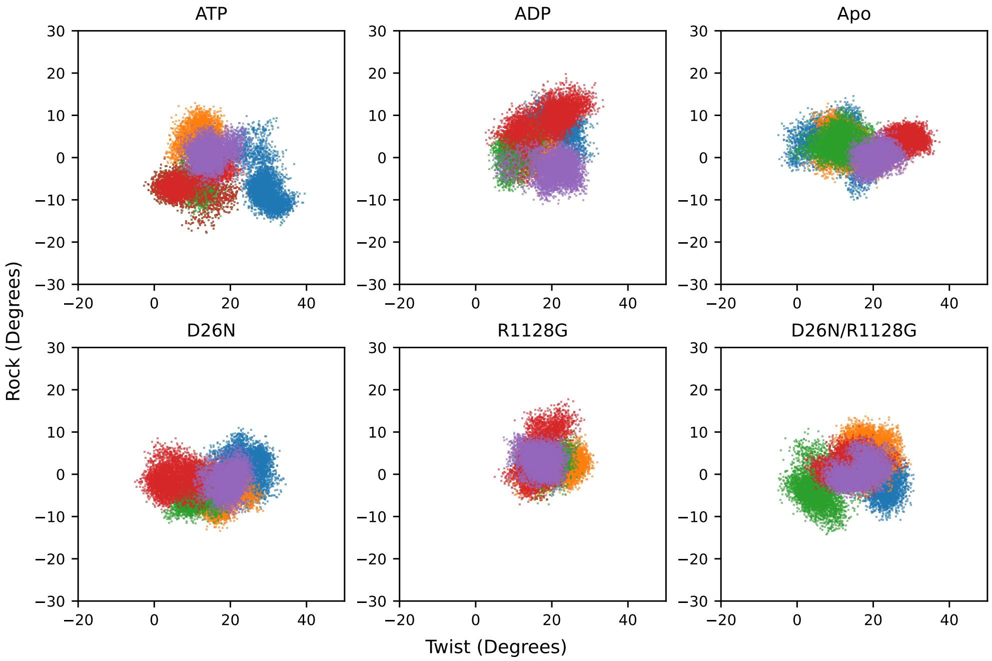


Figure S10: Scatter plots of the twist/rock sampling angles. Each of the five simulations for the six unique systems studies here are colored separately.

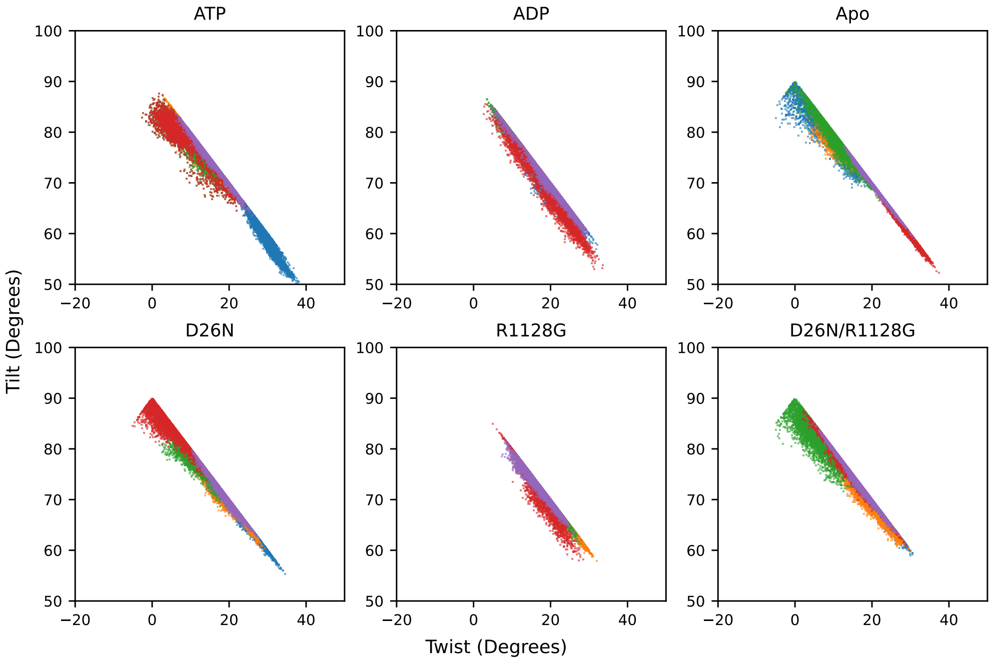


Figure S11: Scatter plots of the twist/tilt sampling angles. Each of the five simulations for the six unique systems studies here are colored separately.

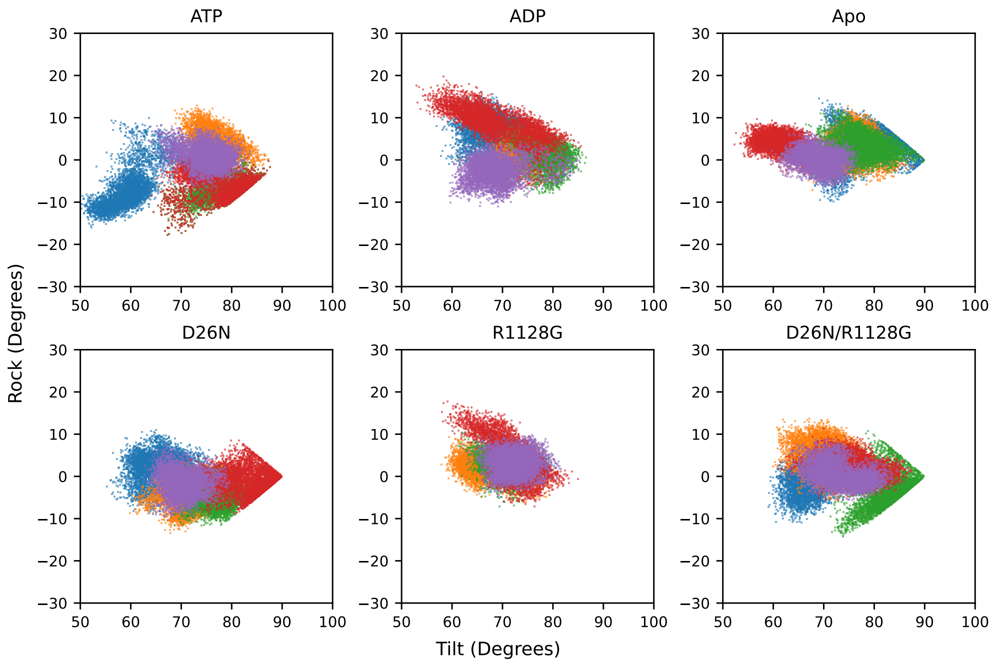


Figure S12: Scatter plots of the tilt/rock sampling angles. Each of the five simulations for the six unique systems studies here are colored separately.

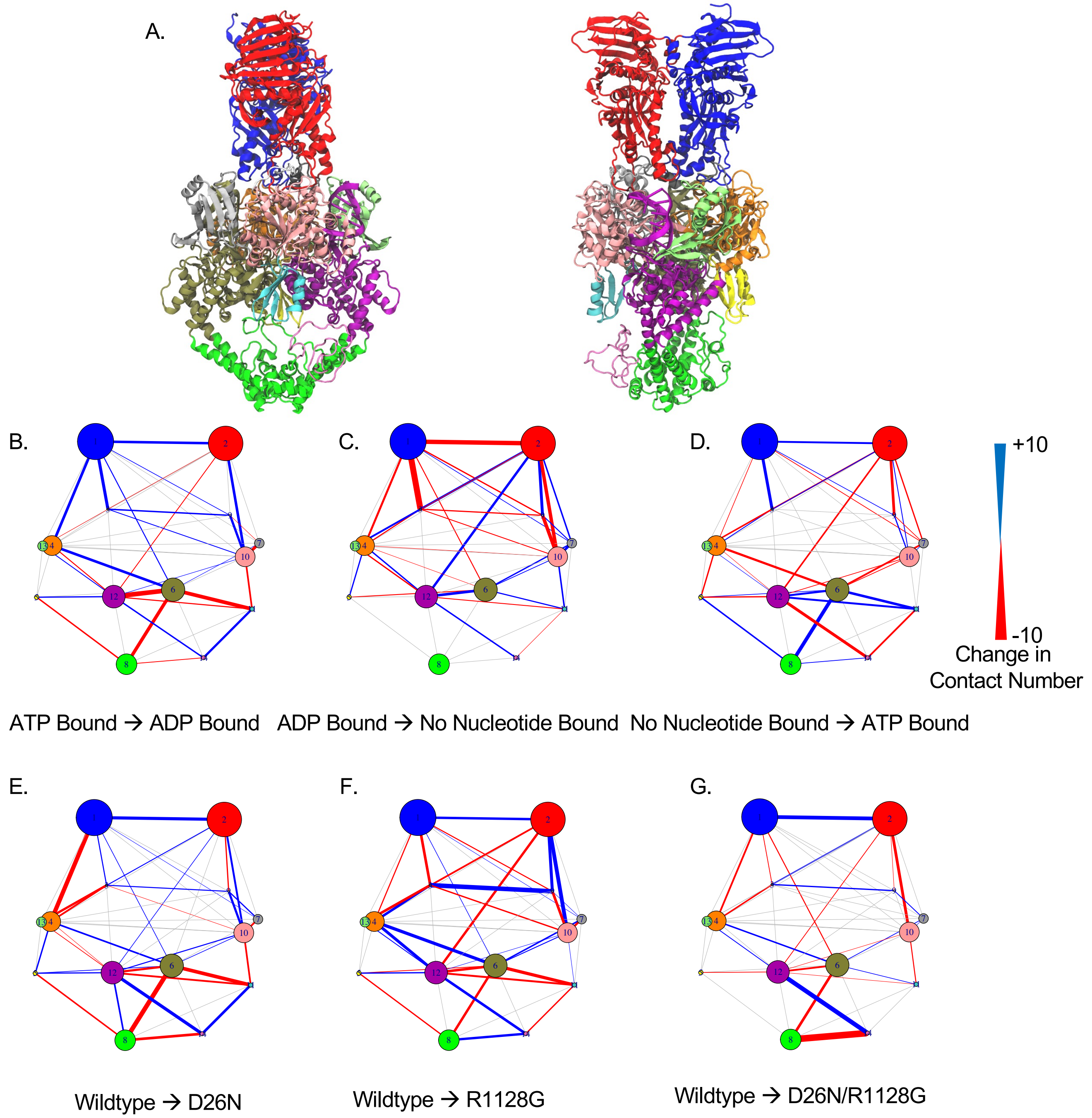


Figure S13: Difference contact network analysis (dCNA) results. **A.** The topoII structure was partitioned into 14 dynamic communities which roughly correspond to complete or partitions of the standard TopoII subdomain assignments ((Figure 1) for the nucleotide binding cycle from ATP to ADP bound (**B.**), ADP to no nucleotide bound (**C.**), and no nucleotide to ATP bound cases (**D.**). dCNA maps were also computed for differences between the ATP bound wildtype and D26N (**E.**), R1128G (**F.**), and D26N/R1128G double mutant (**G.**) states.

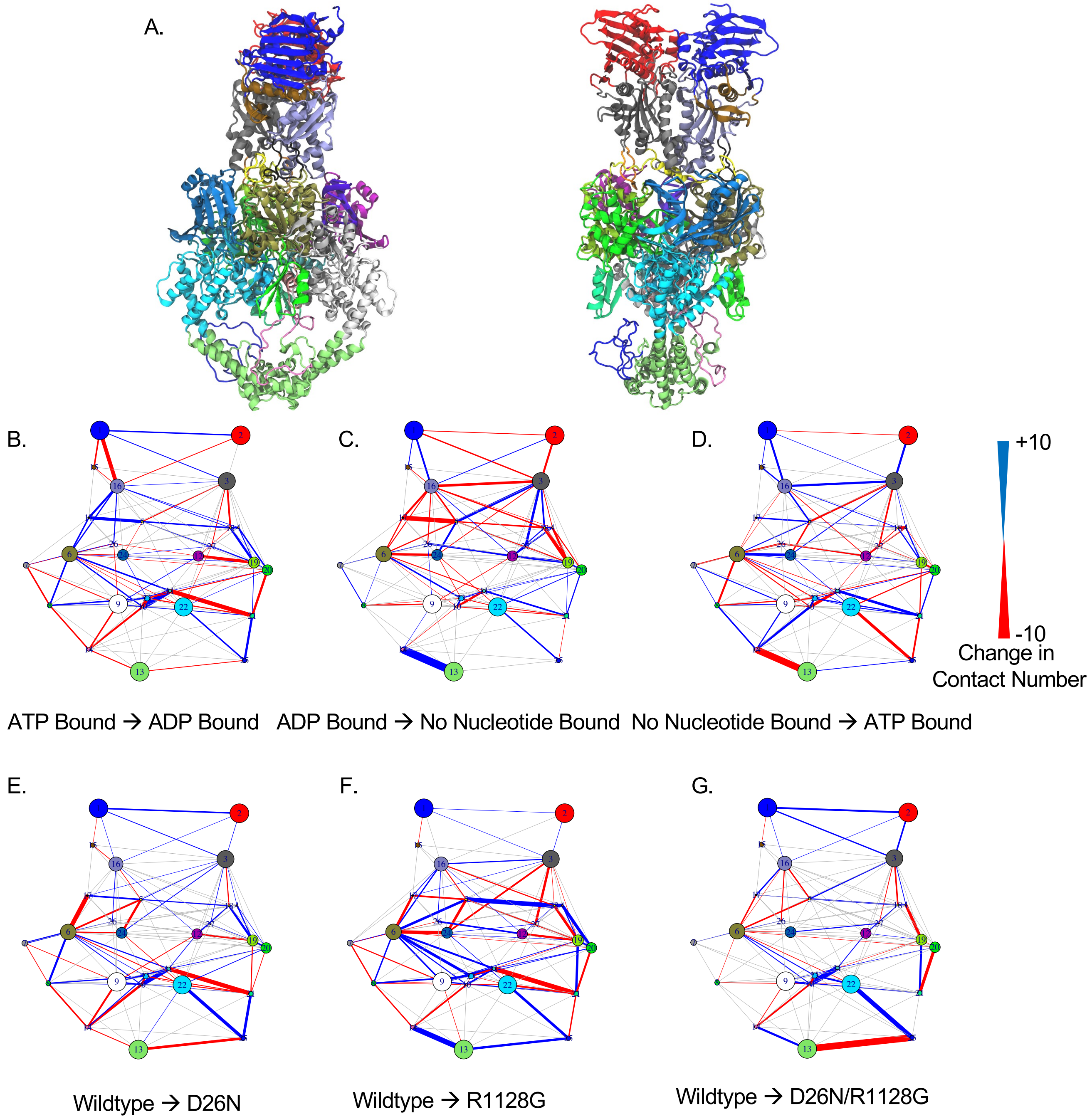


Figure S14: Difference contact network analysis (dCNA) results. **A.** The topoII structure was partitioned into 26 dynamic communities which roughly correspond to complete or partitions of the standard TopoII subdomain assignments. ((Figure 1) for the nucleotide binding cycle from ATP to ADP bound (**B.**), ADP to no nucleotide bound (**C.**), and no nucleotide to ATP bound cases (**D.**). dCNA maps were also computed for differences between the ATP bound wildtype and D26N (**E.**), R1128G (**F.**), and D26N/R1128G double mutant (**G.**) states.

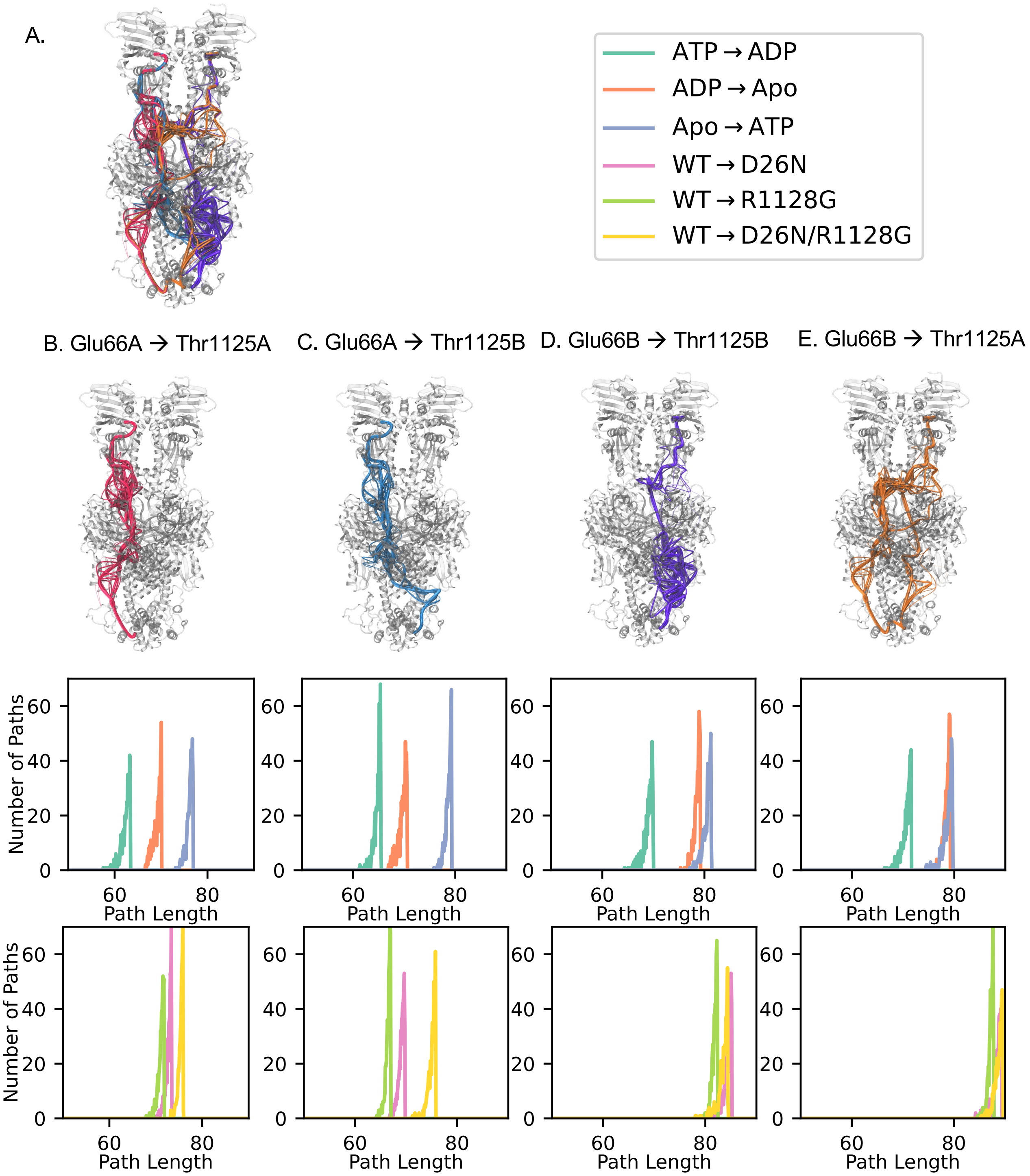


Figure S15: Residue suboptimal pathway analysis for networks beginning at each Glu66 residue and terminating at each Thr1125. (a) Global view of each set of pathways, along with color definitions for pathway length histograms. Pathways for (b) Glu66A→Thr1125A, (c) Glu66A→Thr1125B, (d) Glu66B→Thr1125B, and (e) Glu66B→Thr1125A, along with their respective lengths.

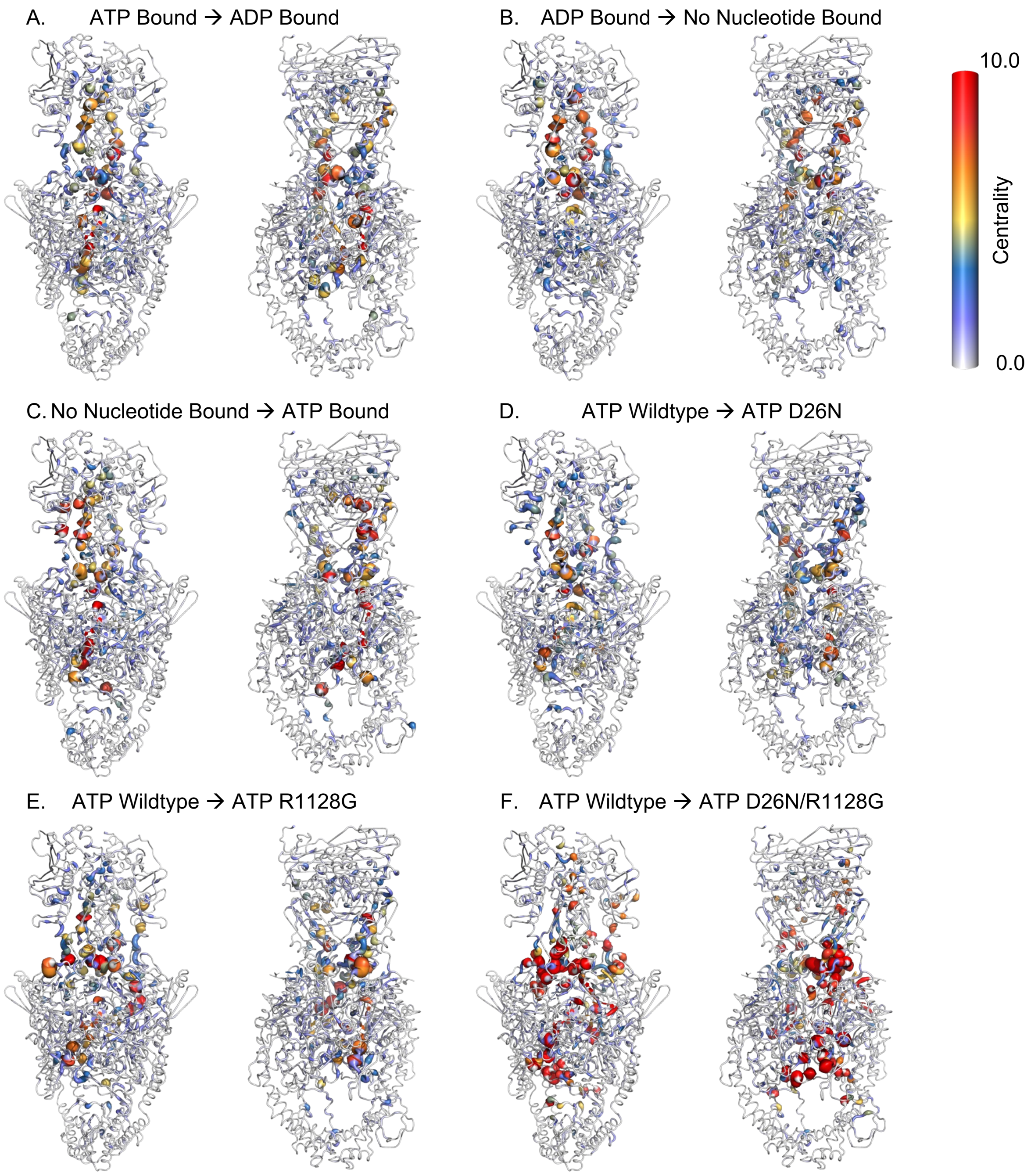


Figure S16: Betweenness centrality scores for all systems.

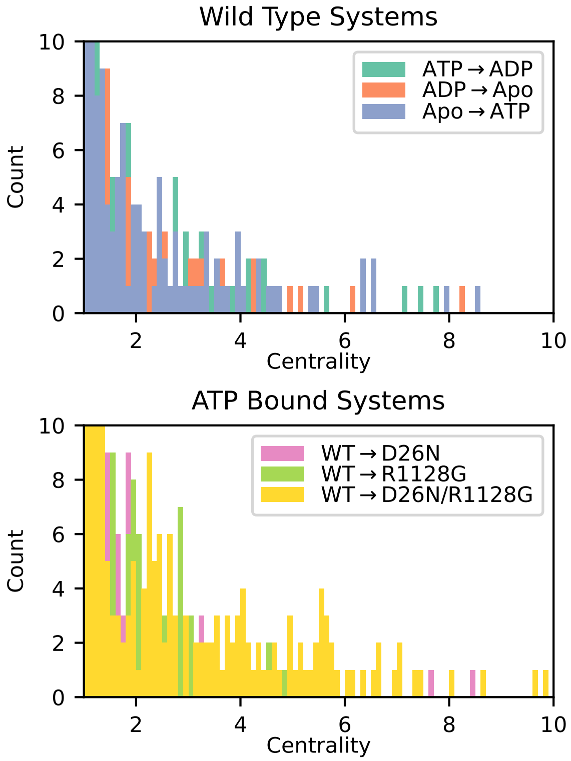


Figure S17: Histogram of betweenness centrality scores for all systems.

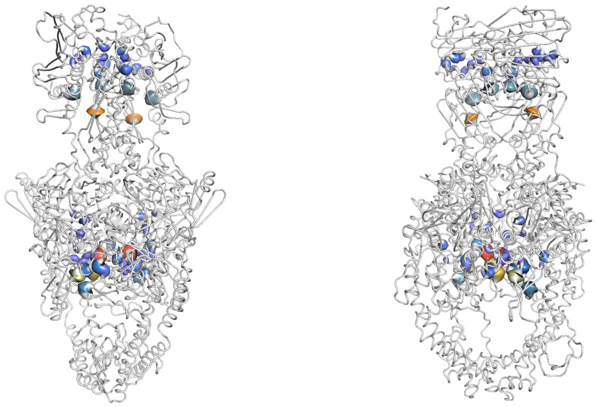

Figure S18: Highly conserved residues found within the top 10 percent of betweenness centrality scores when ordered in descending order. Maximum betweenness centrality score out of three wild type systems were used for each residue. Residues are colored based on the betweenness centrality scores averaged on both monomers.

| Residue Number | Residue | Betweenness centrality Score |
|---|---|---|
| 781 | ARG | 7.900 |
| 352 | PHE | 6.365 |
| 720 | LYS | 5.455 |
| 601 | TYR | 4.580 |
| 225 | ASP | 3.955 |
| 368 | GLU | 3.915 |
| 809 | ASP | 3.760 |
| 605 | LEU | 3.330 |
| 735 | HIE | 3.230 |
| 26 | ASP | 2.800 |
| 690 | ARG | 2.620 |
| 132 | ASP | 2.520 |
| 785 | THR | 2.515 |
| 721 | VAL | 2.490 |
| 77 | ARG | 2.420 |
| 474 | LEU | 2.170 |
| 24 | ARG | 2.160 |
| 734 | TYR | 2.110 |
| 451 | ASP | 2.105 |
| 129 | ASN | 2.020 |
| 71 | ALA | 1.855 |
| 783 | ILE | 1.825 |
| 141 | ARG | 1.760 |
| 603 | LYS | 1.660 |
| 607 | THR | 1.635 |
| 428 | LYS | 1.620 |
| 782 | TYR | 1.475 |
| 74 | ASN | 1.440 |
| 449 | GLU | 1.435 |
| 767 | PHE | 1.415 |
| 724 | LEU | 1.395 |
| 700 | LYS | 1.390 |

Table S1: Highly conserved residues found within the top 10 percent of betweenness centrality scores when ordered in descending order. Maximum betweenness centrality score for each residues out of three wild type systems were used.
